# Temporal transitions in post-mitotic neurons throughout the *C. elegans* nervous system

**DOI:** 10.1101/2020.10.06.328880

**Authors:** HaoSheng Sun, Oliver Hobert

**Affiliations:** Department of Biological Sciences, Howard Hughes Medical Institute, Columbia University, New York, NY, 10027, USA

## Abstract

In most animals, the majority of the nervous system is generated and assembled into neuronal circuits during embryonic development. However, during juvenile stages, nervous systems still undergo extensive anatomical and functional changes to eventually form a fully mature nervous system by the adult stage. The molecular changes in post-mitotic neurons across post-embryonic development and the genetic programs that control these temporal transitions are not well understood. Using the model organism *C. elegans*, we comprehensively characterized the distinct functional states (locomotor behavior) and corresponding distinct molecular states (transcriptome) of the post-mitotic nervous system across temporal transitions from early post-embryonic periods to adulthood. We observed pervasive changes in gene expression, many of which are controlled by the developmental upregulation of the conserved heterochronic miRNA *lin-4/mir-125* and the subsequent promotion of a mature neuronal transcriptional program through the repression of its target, the transcription factor *lin-14*. The functional relevance of these molecular transitions are exemplified by a temporally regulated target gene of the *lin-14* transcription factor, *nlp-45*, a neuropeptide-encoding gene. We found that *nlp-45* is required for temporal transitions in exploratory activity across larval stages, across sexual maturation, and into a diapause arrest stage. Our studies provide new insights into regulatory strategies that control neuron-type specific gene batteries to modulate distinct behaviors states across temporal, sex and environmental dimensions of post-embryonic development, and also provides a rich atlas of post-embryonic molecular changes to uncover additional regulatory mechanisms.

## INTRODUCTION

In non-metamorphosing invertebrates and vertebrates, including mammals, the vast majority of neurons of the adult nervous system are born during embryogenesis. Embryonically born neurons undergo developmental processes, from neuronal identity specification to neuronal migration, axonal guidance/outgrowth and synaptogenesis to form a functional, yet immature nervous system by the time of birth/hatch (Cadwell et al., 2019). During postembryonic stages of life, juvenile nervous systems undergo substantial maturation events that have mostly been characterized on the anatomical and electrophysiological level (Gogtay et al., 2004; Okaty et al., 2009). However, there have been few efforts to systematically characterize the molecular changes within post-mitotic neurons during postembryonic development (Bakken et al., 2016; Kang et al., 2011). It also remains unclear whether post-embryonic, post-mitotic maturation of neurons is mostly a reflection of neuronal activity changes (Spitzer, 2006; Stroud et al., 2020) or whether there are precisely controlled, activity-independent temporal transitions that are genetically hardwired. Genetic programs that control cell fate transitions have been well characterized at earlier stages of neuronal development, on the level of specification of the temporal identity of dividing neuroblasts (neuronal precursor cells), in both vertebrates and *Drosophila* (Cepko, 1999; Holguera and Desplan, 2018; Miyares and Lee, 2019; Pearson and Doe, 2004). However, genetic programs that may specify temporal transitions in post-mitotic neurons during post-embryonic development have been much less considered.

*C. elegans* provides an ideal platform to address these questions with its well-defined post-embryonic life cycle and a well-characterized nervous system with a complete cellular lineage. As with its vertebrate counterparts, the overwhelming majority of the neuron classes present in the full mature adult, hermaphrodite nervous system (97 out of 118 neuron classes) are born, differentiated and synaptically wired during embryogenesis. Hence, young larvae display a functional nervous system capable of executing many sensory and locomotor behaviors that allow the animals to navigate and survive for many weeks or months when developmentally arrested in a diapause state (Baugh, 2013). Some changes in neuronal connectivity, electrophysiology, and behavior during temporal transitions from early larval stages to the adult have been described (Faumont et al., 2006; Fujiwara et al., 2016; Stern et al., 2017; Vidal et al., 2018; White et al., 1978; Witvliet et al., 2020), with the molecular characterization being especially sparse and often without cell-type specific resolution (Boeck et al., 2016; Spencer et al., 2011). The best characterized example of a temporal transition in the *C. elegans* nervous system is the rewiring of the embryonically generated DD motor neurons, a synaptic connectivity change mediated by specific transcription factors, miRNAs and synaptic organizer molecules (Hallam and Jin, 1998; He et al., 2015; Howell et al., 2015; Petersen et al., 2011; White et al., 1978). Here, we ask how broadly such transitions of neuronal features can be found throughout the entire nervous system. Through an analysis of locomotor behavior, controlled by a wide range of neurons throughout the nervous system, we first describe that during larval development, many features of locomotor behavior, including exploratory behavior, are dynamic throughout all post-embryonic temporal transitions. We then provide a panoramic view of gene expression changes throughout the nervous system, and show that many, but not all, of these temporal changes are controlled by the heterochronic pathway (Ambros and Horvitz, 1984; Rougvie and Moss, 2013; Slack and Ruvkun, 1997), specifically the conserved microRNA *lin-4/mir-125* and its downstream effector, the transcription factor *lin-14*. Honing in one of the genes that is temporally controlled by the *lin-4*/*lin-14* axis, the neuropeptide-encoding gene, *nlp-45*, we show that this gene controls temporal transitions in exploratory behavior across larval stages, across sexual maturation, and into a diapause arrest stage. We reveal that the spatiotemporal specificity of *nlp-45* expression across development is controlled in an intersectional manner that integrates temporal (*lin-4*), sexual (TRA-1 master regulator of sex determination), and environmental (insulin-dependent *daf-16*/FOXO) inputs on the level of regulation of *lin-14* transcription factor, with spatial inputs from neuron-type specific, genetically-hardwired terminal selector transcription factors. Together, we present one perhaps generalizable regulatory strategy for dynamic spatiotemporal nervous system gene expression and behavioral transitions across post-embryonic development.

## RESULTS

### Developmental transitions in behavior

To systematically characterize potential changes in nervous system function during post-embryonic neuronal development, we profiled locomotor behavior across all four larval and the adult stage of hermaphrodite *C. elegans* using an automated, high resolution worm tracking system (Yemini et al., 2013). 726 locomotor parameters were separated broadly into 4 categories: morphology, posture, motion, and path. We noticed extensive locomotor behavioral changes in all 4 categories across all post-embryonic temporal transitions (**Fig.1A**). While some of the motion and path features might be attributed to developmental changes in non-neuronal tissues such as muscle, many of the developmentally regulated motion and path features were likely to be mediated exclusively by differences in nervous system function across development (**Fig.1A**). For example, animals at the second larval (L2) stage exhibited increased pausing and dwelling behavior compared to animals at the first larval (L1) stage while adult animals exhibited increased forward motion and decreased backward motion compared to animals at the last larval stage (L4) (**Fig.1A**). Some of these specific changes, such as dwelling and pausing, have been previously used to define distinct states in exploratory behavior (Flavell et al., 2013; Fujiwara et al., 2002).

**Figure 1:**
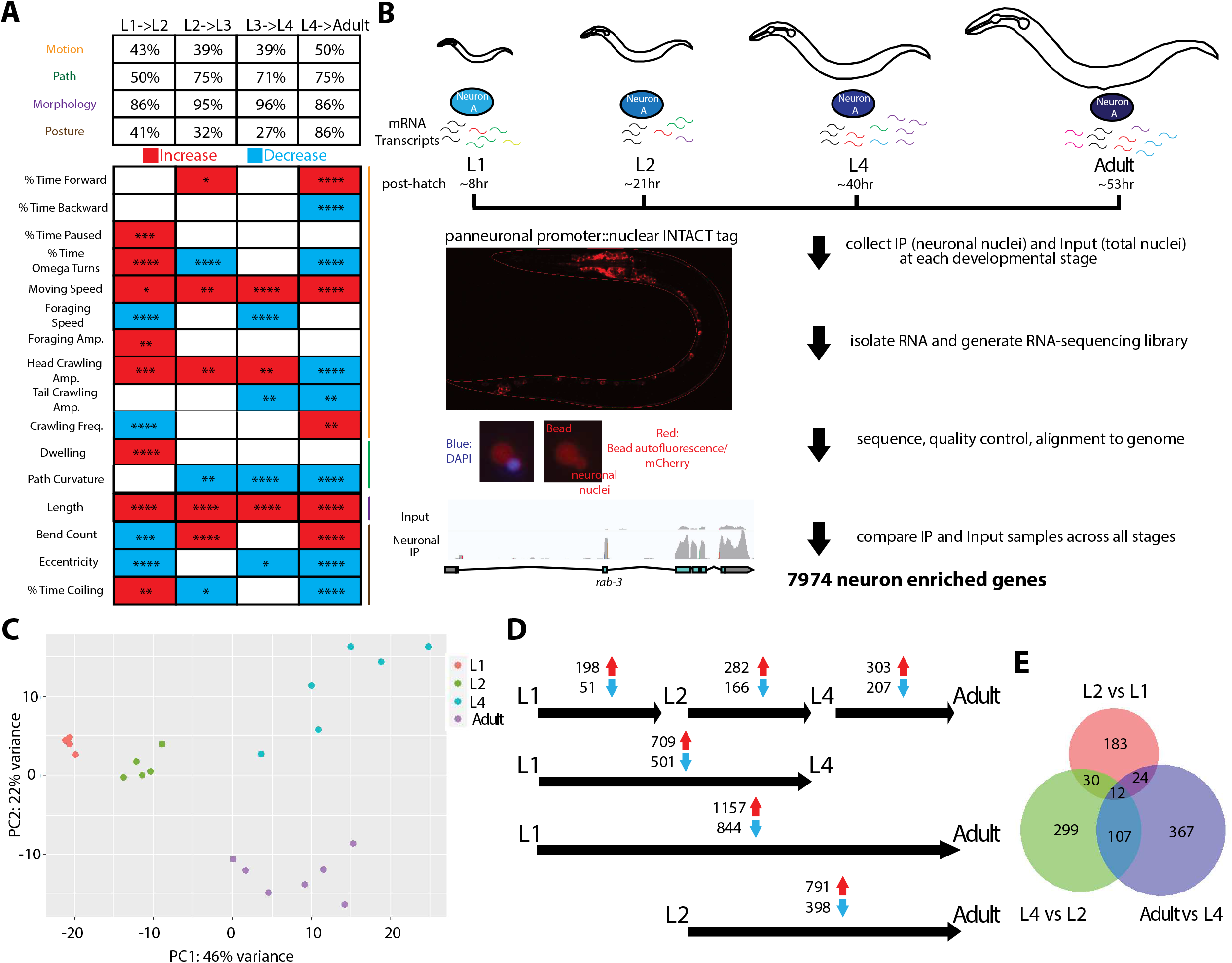
Temporal transitions in behavior and neuronal transcriptome across *C. elegans* post-embryonic development. **(A)** Developmental transition in locomotor behavior across post-embryonic life stages, as measured by automated single worm tracking (Yemini et al., 2013). The upper panel shows the percentage of parameters in each of the 4 broad categories that are different across each transition (q<0.05). The bottom panel shows representative parameters from each category, as indicated by color-coded lines on the right, across development. Red and blue rectangles represent increases and decrease in each parameter, respectively, across the specified transitions. Number of animals tracked for each stage are as follow: L1,47; L2,48; L3,41; L4,129; Adult,107. Wilcoxon rank-sum tests and false-discovery rate q values for each comparison are presented in each rectangle: *q<0.05; **q<0.01, ***q<0.001, ****q<0.0001. Amp: Amplitude. Freq: Frequency. **(B)** Schematic and experimental design for INTACT sample collection, protocol, and data analysis for neuronal transcriptome profiling across development. Representative images of the panneuronal INTACT strain as well as neuronal nuclei after immunoprecipitation (IP) are shown in bottom left panels. Representative tracks from IGV are shown for input and neuronal IP samples to demonstrate IP enrichment for panneuronally expressed gene, *rab-3*. **(C)** Distinct molecular states (neuronal transcriptome) across post-embryonic developmental stages. Each dot represents a replicate in the RNA-seq analysis. The increased variabilities of L4 and adult samples were as a result of batch effect, and was taken as a variable for downstream analysis. Principal component analysis (PCA) of neuronal transcriptome across post-embryonic development was conducted using DESeq2 in R studio (Love et al., 2014). **(D)** Developmental transitions in neuronal transcriptome across post-embryonic life stages. The numbers of significant (p_adj_<0.01) increases/decreases in gene expression are shown for each stage transition. **(E)** Venn diagram of developmental changes in neuronal gene expression across different stage transitions, showing some overlaps but mostly distinct developmental changes across each stage transition. For all panels, L1 through L4 represent the first through the fourth larval stage animals.

### Transitions in gene expression throughout the nervous system

The extensive changes in locomotor and exploratory behavior (*distinct behavioral states*) across post-embryonic development suggested the existence of changes in neuronal gene expression (*distinct molecular states*) across post-embryonic development (**Fig.1B**). Rather than focusing on specific neurons, we took a panoramic approach and profiled the transcriptome of the entire nervous system using INTACT technology (Deal and Henikoff, 2010; Steiner et al., 2012). To this end, a panneuronal promoter was used to express a FLAG-tagged nucleoporin protein (NPP-9) for immunoprecipitation of neuronal nuclei, at 4 post-embryonic stages: 3 larval stages (L1, L2, and L4) and the adult (8 +/− 2 hours after the L4/adult molt) stage (**Fig.1B**). After comparing the transcriptome of immunoprecipitated (IP) neuronal nuclei samples to input samples (total nuclei), we identified 7974 genes that were neuronally-enriched (p_adj_ <0.05) and proceeded with all downstream analysis focusing on these genes (**Table S1, S2**). In addition to recapitulating previously described neuronally-enriched genes (Kaletsky et al., 2016; Spencer et al., 2011), our panneuronal profiling was able to capture all neuronal classes, as the top uniquely expressed genes in each neuronal class were well represented in our panneuronal IP data (Taylor et al., 2019). A principal-component analysis (PCA) on the expression profile of these neuronally-enriched genes across post-embryonic developmental stages revealed that the neuronal transcriptome of each developmental stage clustered together and were distinct from the other stages, suggesting developmental stage-specific molecular states for the nervous system (**Fig.1C**). Comparison of differential gene expression revealed both up and downregulation across all developmental transitions (**Fig.1D**), with some genes being dynamic across all temporal transitions while many genes demonstrated transcriptomic changes only at specific stage transitions (**Fig.1E**). All in all, 2639 genes (p_adj_ < 0.01; ~33% out of the 7974 neuronally-enriched genes) exhibited transcriptomic changes at some point during post-embryonic development (**Table S3, S4**). Together, this data demonstrated that extensive molecular changes were occurring during post-mitotic post-embryonic neuronal development.

We observed three major patterns of gene expression transitions among these 2639 developmentally-regulated neuronally-enriched genes (**Fig.2A**): 1) cohorts of genes whose expression decreased across development, particularly from early larval to mid/late larval stages; 2) other cohorts of genes whose expression increased across development, particularly during transition from early/mid larval stages to late larval stage/adulthood; 3) additional cohorts of genes whose expression peaked at the last larval stage and subsequently decreased upon transition into adulthood. Due to the panneuronal nature of our profiling, such increases or decreases in gene expression could entail binary on/off switches in individual neuron types and/or could entail relative changes in levels of expression in the same set of neurons. Using gene expression reporters that detect changes with single neuron resolution (many of them endogenous reporter alleles that we engineered with CRISPR/Cas9 technology), we found ample evidence for both scenarios (**Fig.2, S1**). For example, expression of the glutamate receptor *gbb-2* decreased in all expressing cells throughout larval development (**Fig.2B**), while expression of the innexin *inx-19* was progressively lost from specific neuron types throughout larval stages (**Fig.2C**). Moreover, we detected uniform changes of broadly expressed genes and, on the opposite end of the spectrum, changes of very restrictively expressed genes with up-or down-regulation in small subset of neurons or even a single neuron class. For example, the transcriptional cofactor MAB-10 was undetectable in early larval stages, and its expression across the entire nervous system was turned on at the L4 stage and further upregulated in adulthood (**Fig.2D**). On the other end of the spectrum, the insulin-related neuropeptide *ins-6*, was only expressed in the ASI sensory neurons in L1 and L2 stage animals, and its expression was turned on in 1 additional pair of neurons, ASJ, during the L2 to L3 transition and maintained into adulthood (**Fig.2E**). Similarly, while the neuropeptide *nlp-13* remained stable in a subset of pharyngeal neurons, it became upregulated specifically in the IL2 neurons (**Fig.2G**). As another example, expression of *ins-9* switched on in a number of neurons at the L2 to L3 transition, and subsequently lost expression in a subset of these neurons upon transition into adulthood (**Fig.2F**). Additional validated changes are documented in **Fig.S1**. The neurons that demonstrated developmentally regulated changes spanned all type (sensory vs inter-vs motor-neurons), neurotransmitter usage (glutamate vs acetylcholine vs GABA vs dopamine), and lineage history (**Fig.2, S1**)

**Figure 2:**
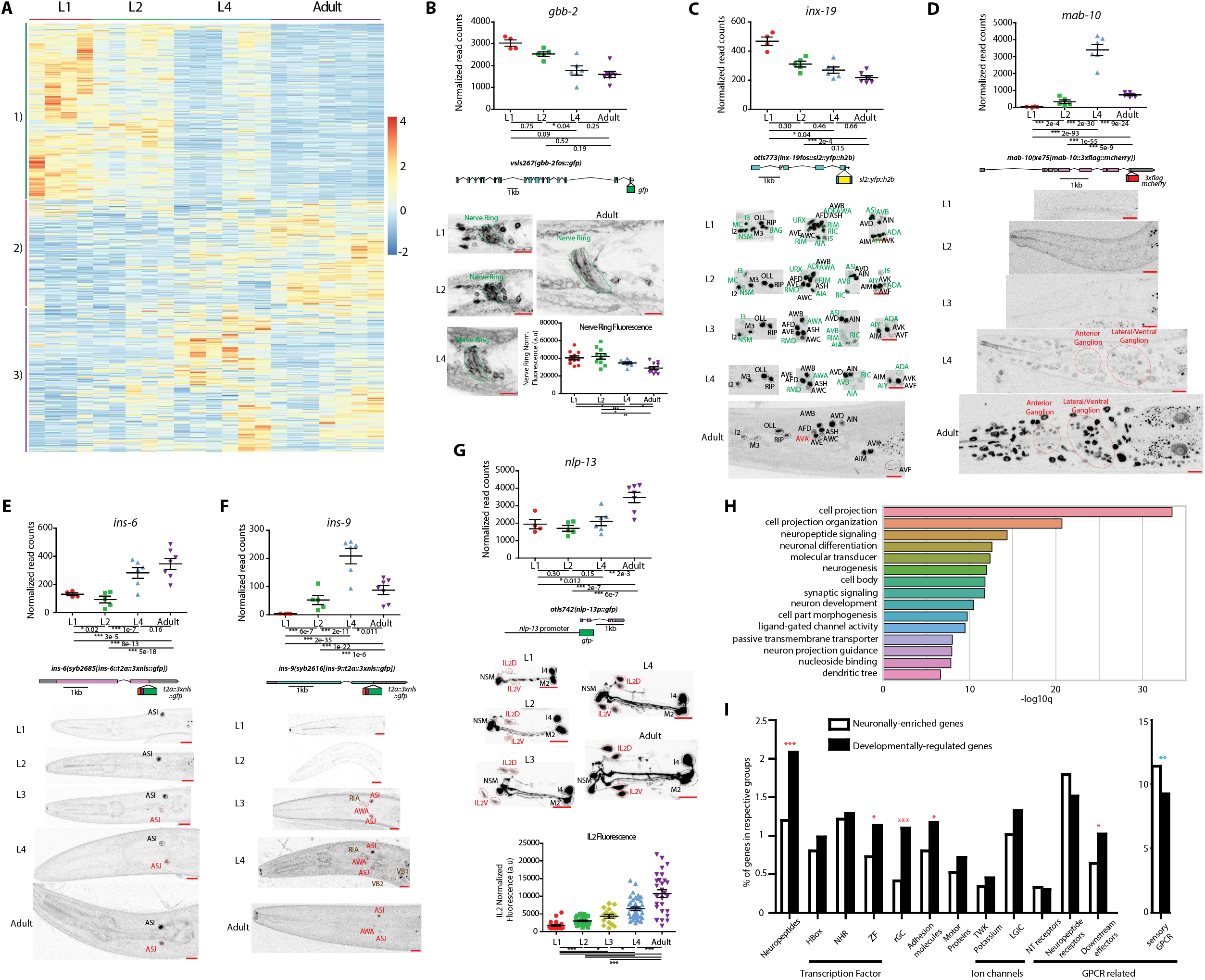
Temporal transitions in nervous system gene expression across *C. elegans* post-embryonic development. **(A)** Heatmap of the 2639 developmentally regulated genes (p_adj_<0.01) across post-embryonic life stages. Values were z-score normalized and plotted using pheatmap in R studio. Each row represents a single gene, and each column represents a single RNA-seq replicate. The three major patterns of developmental changes in neuronal gene expression, as discussed in the text, are indicated by colored lines and numbers on the left side. Briefly, 1), 2), and 3) represents cohorts of genes whose expression decreased across development, increased across development, and peaked at the last larval stage before decreasing in adulthood, respectively. The increased variabilities of L4 and adult samples were as a result of batch effect, and was taken as a variable for downstream analysis. For (B)-(G), validations of developmentally regulated genes with expression reporters are shown. On top are the scattered dot plots (each point represents a single replicate) of the normalized read counts across all developmental stages from the neuronal INTACT/RNA-seq profiling. Mean +/− SEM are shown for each stage. Adjusted p values (p_adj_) for each developmental comparison are below. For p_adj_, *<0.05, **<0.01, ***<0.001. Below the RNA-seq read count plots are the schematics and allele names of the expression reporters. Below that are representative confocal microscopy images of the expression reporters across development. Specific regions/neurons are labeled with dotted lines: those labelled with black dotted lines/names are not altered developmentally while those labelled with green and red lines/names demonstrate, respectively, decreases and increases in expression across development. Those labelled with brown lines/name demonstrate both increases and decreases in expression in the same neurons across development. For (B) and (G), additional quantifications of fluorescence intensity are also shown at the bottom. Red scale bars (10μm) are on the bottom right of all representative images. **(B)** Metabotropic glutamate receptor *gbb-2*, as validated with a translational fosmid reporter (*gfp*), shows expression in the same set of neurons across development (Yemini et al., 2019), although the intensity of expression is decreased across development, including that in the nerve ring as measured with fluorescence intensity. **(C)** Gap junction molecule *inx-19*, as validated with a transcriptional fosmid reporter (*sl2::yfp::h2b*), loses expression in sixteen neuronal classes across development and gains expression in the AVA neuron upon entry into adulthood. **(D)** Transcription cofactor *mab-10*, as validated with an endogenous translational reporter (*3xflag::mcherry*) engineered with CRISPR/Cas9, gains expression across the nervous system amongst other tissue during transition into the L4 stage that is further upregulated in adulthood. The RNA prediction matches well with previous RNA FISH analysis (Harris and Horvitz, 2011). The difference between RNA data and protein reporter expression is consistent with previous characterized post-transcriptional regulation by LIN-41 (Aeschimann et al., 2017). **(E)** Neuropeptide-encoding gene *ins-6*, as validated with an endogenous reporter (*t2a::3xnls::gfp*) engineered with CRISPR-Cas9, gains expression in ASJ across the L2->L3 transition. **(F)** Neuropeptide-encoding gene *ins-9*, as validated with an endogenous reporter (*t2a::3xnls::gfp*) engineered with CRISPR-Cas9, gains expression in a number of neurons as it enters L3/L4 stages and loses expression in a subset of these neurons upon entry into adulthood. **(G)** Neuropeptide-encoding gene *nlp-13*, as validated with a promoter fusion (*gfp*) reporter, maintains expression in a number of pharyngeal neurons while gaining expression (as measured by fluorescence intensity) in the IL2 neurons across development. **(H)** Gene ontology analysis of the 2639 developmentally-regulated genes using the Enrichment Tool from Wormbase. **(I)** Enrichment or depletion of gene families in the subset of 2639 developmentally regulated genes in comparison to all 7974 neuronal enriched genes. The z score test for two population proportion is used to calculate statistically significance of the comparisons. *p<0.05; **p<0.01, ***p<0.001. Red and blue * indicates over-representation and under-representation, respectively, of the gene family in the developmentally-regulated gene battery in comparison to all neuronal enriched genes. HBox: homeodomain transcription factors; NHR: nuclear hormone receptor transcription factors; ZF: zinc finger transcription factors, rGC: receptor-type gunacyl cyclase; LGIC: ligand-gated ion channels; NT: neurotransmitter; GPCR: G-protein coupled receptors. For all panels, L1 through L4 represent the first through the fourth larval stage animals. Additional changes are documented in Fig.S1.

Gene ontology (GO) analysis of the developmentally regulated genes revealed an expected enrichment in nervous system-associated genes, including neuronal projection/guidance, neuropeptide/synaptic signaling, and ligand-gated channel activity (**Fig.2H**). Specific gene families were over-represented amongst the developmentally regulated genes, including neuropeptides, receptor-type guanylyl cyclases (rGCs), cell adhesion molecules, GPCR-associated downstream effectors and zinc-finger transcription factors; while GPCRs were slighted under-represented (**Fig.2I**). Within individual gene families, many members displayed temporal changes in global gene expression patterns, best exemplified with rGC sensory receptors and neuropeptide-encoding genes (**Fig.S1A-G**).

### Many, but not all temporal transitions are regulated by the *lin-4/lin-14*-initiated heterochronic pathway

To investigate the regulation of these temporal transitions, we considered the heterochronic pathway, a cascade of microRNAs, RNA-binding proteins, and transcription factors that had been shown to regulate the temporal developmental progression in mitotic ectodermal lineages and reproductive system (Ambros and Horvitz, 1984; Rougvie and Moss, 2013; Slack and Ruvkun, 1997) (**Fig.3A**), as well as some temporal transitions in post-mitotic nervous system such as DD rewiring and the manifestation of sexually dimorphic phenotypes (Hallam and Jin, 1998; Howell et al., 2015; Lawson et al., 2019; Pereira et al., 2019). The conserved microRNA *lin-4*/*mir-125* marks the first transition in this pathway and through inhibiting its targets, triggers downstream events in the pathways. Specifically, *lin-4* upregulation, as determined by whole animal Northern Blots, downregulates its downstream target, the transcription factor *lin-14* at the L1 to L2 transition (Feinbaum and Ambros, 1999; Ruvkun and Giusto, 1989) (**Fig.3A**).

**Figure 3:**
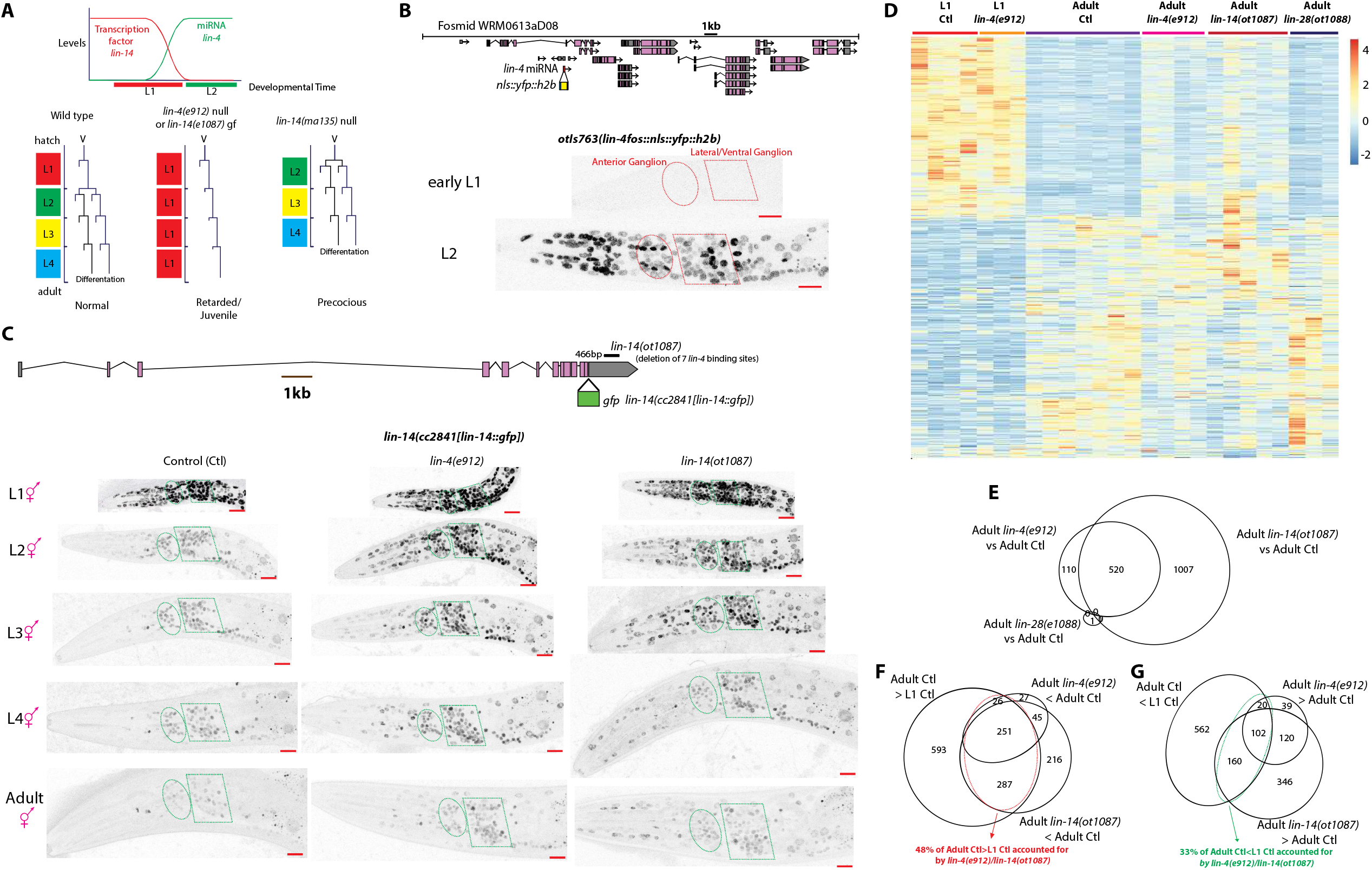
MicroRNA *lin-4* mir-125 partially juvenizes the neuronal transcriptome through regulation of the transcription factor *lin-14*. **(A)** Heterochronic pathway in mitotic epithelial/hypodermal lineages. Upper panel is a simplified schematic that shows the upregulation of *lin-4* miRNA and the subsequent repression of transcription factor *lin-14* during the L1 to L2 transition. Bottom panel shows the heterochronic phenotype of *lin-4* and *lin-14* mutants in a representative epithelial/hypodermal (V) lineage. Compared to wild type, the V lineage of *lin-4* null and *lin-14* gain of function(gf) mutants shows retarded/juvenile pattern of mitotic divisions, reiterating L1-like divisions throughout development, while the V lineage of *lin-14* null mutants shows precocious L2-like mitotic division patterns at the L1 stage and differentiate one stage earlier. The content/formatting of the bottom panel is adapted from a previous review (Moss, 2007). **(B)** Expression of *lin-4* is turned on in the nervous system amongst other tissue during the L1->L2 transition. Schematic of the *lin-4* fosmid expression reagent is shown above. Ellipse and polygon outlines the anterior and lateral/ventral neuronal ganglions respectively. **(C)** Expression of *lin-14* is downregulated in the nervous system amongst other tissue across post-embryonic development in a *lin-4* dependent manner in hermaphrodite animals. Schematic of the *lin-14* translational GFP allele, as engineered by CRISPR-Cas9, as well as a *lin-14(ot1087)* gain of function allele, where a 466bp region containing all seven *lin-4* repressive binding sites is deleted, is shown above. Ellipse and polygon outlines the anterior and lateral/ventral neuronal ganglions respectively. Expression of *lin-14* is still detectable in the adult hermaphrodite. *lin-14* expression is upregulated in *lin-4(e912)* null and *lin-14(ot1087)* gain of function animals across development. **(D)** *lin-4* null mutation partially juvenizes the adult control(Ctl) neuronal transcriptome to resemble that of the L1 Ctl neuronal transcriptome through de-repression of *lin-14* and not *lin-28*. The *lin-14(ot1087)* gain of function and *lin-28(ot1088)* gain of function alleles were generated through deletion of *lin-4* binding sites in their respective 3’UTR (Fig. 3C and S2). Values were z-score normalized and plotted using pheatmap in R studio. Each row represents a single gene, and each column represents a single RNA-seq replicate. **(E)** Venn diagram showing that the difference between the adult *lin-4* null neuronal transcriptome compared to the adult control(Ctl) neuronal transcriptome is largely recapitulated in the transcriptome of adult *lin-14(ot1087)* gain of function mutants. Only one gene is significantly different in the adult *lin-28(ot1088)* gain of function vs adult control comparison and does not overlap with the genes regulated by *lin-4/lin-14*. **(F)** Venn diagram showing that 48% of genes that demonstrate developmental upregulation (adult control(Ctl)>L1 Ctl) are at least partially juvenized in the adult *lin-4* null and/or *lin-14(ot1087)* gain of function animals. **(G)** Venn diagram showing that 33% of genes that demonstrate developmental downregulation (adult control(Ctl)<L1 Ctl) are at least partially juvenized in the adult *lin-4* null and/or *lin-14(ot1087)* gain of function animals. For panels (B) and (C), L1 through L4 represent the first through the fourth larval stage animals. Red scale bars (10μm) are on the bottom right of all representative images.

Since *lin-4/mir-125* and *lin-14* expression have not been precisely examined throughout the nervous system, we analyzed the expression of both genes using either fosmid based reporter genes and/or CRISPR/Cas9-engineered reporter alleles. A fosmid based reporter gene and a CRISPR/Cas9-engineered reporter allele in which the *lin-4* pre-miRNA sequence was replaced with YFP (fosmid) or mScarlet (allele), displayed an upregulation of *lin-4* expression at the L1 to L2 transition in most cells, including the nervous system (**Fig.3B**; data not shown), agreeing with previous studies in whole animal lysates (Bracht et al., 2010; Feinbaum and Ambros, 1999). A CRISPR/Cas9 engineered endogenous GFP reporter of *lin-4’s* downstream target, *lin-14*, revealed *lin-4*-dependent downregulation of LIN-14::GFP throughout the entire nervous system at the L2 stage (**Fig.3C**). Further downregulation of neuronal *lin-14* expression was observed beyond the L2 stage, but LIN-14::GFP was still detectable in the neurons of adult animals (**Fig.3C**).

The patterns of *lin-4* and *lin-14* expression suggested that the *lin-14* transcription factor may promote a juvenile neuronal gene expression pattern, while upregulation of *lin-4* and subsequent downregulation of *lin-14* may be required to promote a mature neuronal gene expression program. To test this hypothesis, we again used INTACT to profile the neuronal transcriptome of *lin-4* null and *lin-14(ot1087)* gain of function adult animals, where all seven *lin-4* repressive binding sites were deleted in the *lin-14* 3’UTR, which resulted in sustained neuronal expression of *lin-14* throughout post-embryonic development and phenocopied the *lin-4* null mutants (**Fig.3C**). The transcriptome profiling revealed that *lin-4* null and *lin-14(ot1087)* gain of function mutations broadly juvenized neuronal transcriptome in adults: 48% and 33% of genes that were upregulated and downregulated, respectively, between the first larval and adult stage were at least partially juvenized by *lin-4* null and/or *lin-14(ot1087)* gain of function mutations (**Fig. 3D-G, Table S5**). The partial nature of the juvenization of the neuronal transcriptome by *lin-4* null and *lin-14(ot1087)* gain of function mutations suggested that there must be additional mechanisms beyond *lin-4* and *lin-14* triggered heterochronic pathway that regulate the temporal transition of the nervous system across development. The other well-characterized downstream effector of *lin-4*, the conserved RNA-binding protein *lin-28*/*LIN-28* (Moss et al., 1997), did not account for any of the juvenizing effect of *lin-4* null mutant on the adult neuronal transcriptome (**Fig.3D, E, Table S5**).

#### Regulation of *nlp-45* by the *lin-4* miRNA and its target, LIN-14

To validate and further explore the impact of *lin-4* and *lin-14* on neuron-specific gene expression patterns, we focused on one neuropeptide, *nlp-45*, that our transcriptome profiling approach identified to be temporally controlled across all examined temporal transitions and regulated by *lin-4*/*lin-14* (**Fig.4A, B**). Our focus on a neuropeptide was motivated by the over-representation of neuropeptides in our developmentally regulated gene battery (**Fig.2I**) and by their well-known function as modulators of transitions between discrete behavioral states (Marder, 2012; Schoofs and Beets, 2013; Waggoner et al., 2000). We first validated and determined the cell type specificity of the temporal regulation of *nlp-45* using a CRISPR/Cas9-engineered reporter allele, in which we inserted a nuclear localized GFP behind the NLP-45 coding sequences, separated by a T2A sequence that splits the two proteins (Ahier and Jarriault, 2014) (**Fig.4A**). At hatching, *nlp-45* was consistently expressed in one pair of interneurons, RIA, with occasional dim expression in the AFD and AWA sensory neurons, remnants of embryonic expression that became invisible by the mid-L1 stage. At the L2 stage, *nlp-45* expression commenced in the head motor neurons RMDD/V, and variably in the OLL and CEPD/V sensory neurons, which became consistently expressed by the L3 stage. From the L3 stage onwards, *nlp-45* expression was also observed in the PVD, ADE, PDE sensory neurons, and variably, in neurons in the RVG. Upon entry into adulthood, additional dim and variable *nlp-45* expression was observed in A- and B-type ventral nerve cord (VNC) motor neurons (**Fig.4A, 6A, E**).

**Figure 4:**
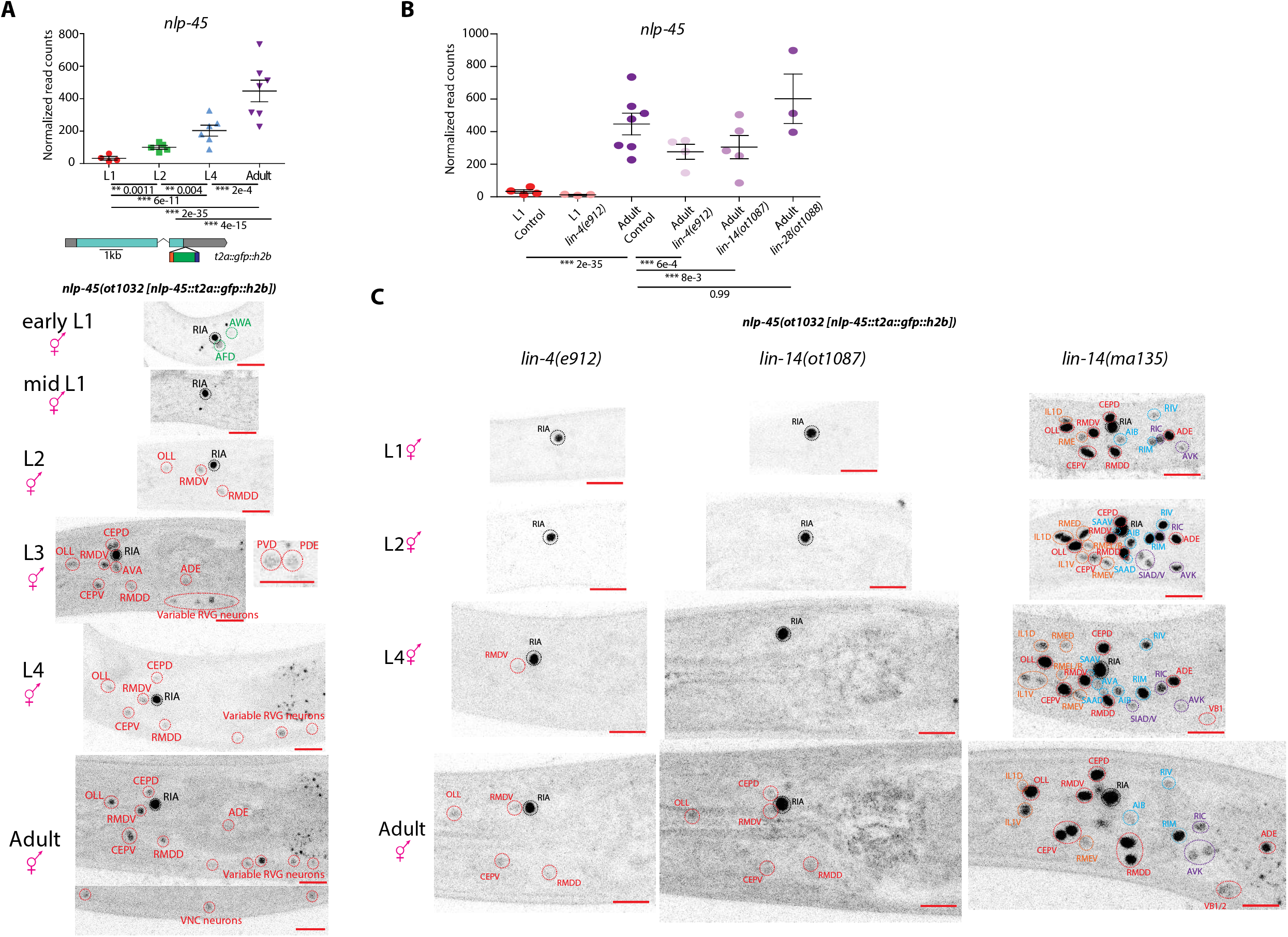
*lin-4/lin-14* controls the temporal expression pattern of neuropeptide-encoding gene, *nlp-45*. **(A)** Temporal expression pattern of *nlp-45*. On top is the scattered dot plot (each point represents a single replicate) of the normalized read counts across all developmental stages from the neuronal INTACT/RNA-seq profiling. Mean +/− SEM are shown for each stage. Adjusted p values (p_adj_) for each developmental comparison are below. For p_adj_, *<0.05, **<0.01, ***<0.001. Below the RNA-seq read count plot are the schematic and allele name of the CRISPR-Cas9 engineered endogenous expression reporter (*t2a::3xnls::gfp*). Below that are representative confocal microscopy images of the expression reporter across development. Specific region/neurons labeled with dotted lines: those labelled with black dotted lines/names are not altered developmentally while those labelled with green and red lines/names demonstrate, respectively, decreases and increases in expression across development. Other than some remnant expression from embryo in early L1 animals, *nlp-45* gains expression progressively in a number of neurons across development. **(B)** Juvenization of *nlp-45* expression by *lin-4*/*lin-14* across development as predicted from the neuronal INTACT/RNA-seq profiling. Normalized RNA-seq read counts for the L1/adult control/mutant animals are plotted, with each point representing a replicate and the mean +/− SEM shown for each stage. ***p_adj_<0.001. **(C)** Expression reporter validation of the effect of *lin-4*/*lin-14* on *nlp-45* expression. *nlp-45* expression is juvenized in *lin-4*(e912) null and *lin-14*(ot1087) gain of function animals (left and middle columns) while precocious *nlp-45* expression is observed in the *lin-14(ma135)* null mutants, where neurons that typically expresses *nlp-45* in later larval/adult hermaphrodite stage (labelled in red), in adult males (labelled in blue), and in dauer animals (labelled in orange) show expression at the L1 stage (right column). Additional neurons that never show expression in any conditions tested (labelled in purple) also show *nlp-45* expression in *lin-14(ma135)* null animals. For all panels, L1 through L4 represent the first through the fourth larval stage animals. Red scale bars (10μm) are on the bottom right of all representative images.

In the *lin-4* null mutant, we observed a “juvenized” pattern of *nlp-45* expression as predicted by our transcriptome profiling experiment: expression remained largely restricted to the RIA interneuron throughout development, with variable and very dim expression in the RMDD/V, OLL, and CEPD/V neurons starting in late L4/adult stage (**Fig.4B, C**). Similarly, in the *lin-14(ot1087)* gain of function mutant, we observed a recapitulation of the juvenized expression pattern as observed in the *lin-4* null mutant (**Fig.4B, C**). Conversely, precocious *nlp-45* expression was observed in the *lin-14(ma135)* null mutant, where cells that typically expressed *nlp-45* at later stages showed expression at the L1 stage (**Fig.4C**). Consistent with the profiling experiments, *lin-28* gain or loss of function mutants did not affect the developmental *nlp-45* expression profile (**Fig.4B, S2**), suggesting that *lin-4* regulated the maturation of the neuronal transcriptome largely through regulating the LIN-14 transcription factor.

#### *nlp-45* modulates temporal transition in exploratory behavior

Given the tightly controlled, *lin-4/14*-dependent expression pattern of *nlp-45* across post-embryonic development, we set out to functionally characterize the role of *nlp-45* across development. One of the more dramatic transitions in *nlp-45* expression in hermaphrodites was the transition between L1 and later larval stages, when expression broadened from RIA to additional sets of neurons (**Fig.4A**). One of the locomotor behavioral parameters modulated during this temporal transition was the increased dwelling behavior of L2 stage animals compared to L1 stage animals, demonstrated in two independent behavioral assays: the automated worm tracking and an adapted exploratory assay (Flavell et al., 2013; Yemini et al., 2013) (**Fig.1A, 5A, B, C**). Consistent with its broadened expression pattern at later larval stages, two different deletion alleles of *nlp-45*, generated by CRISPR-Cas9, resulted in a stage specific increase in exploratory behavior only observed for L2, and not L1, animals (**Fig. 5A**). Transgenic expression of *nlp-45* in either the RMDD/V or RIA neurons reversed the increased exploratory behavior of L2 *nlp-45* mutants to that of L2 control (N2) animals (**Fig.5A**). The ectopic/overexpression of *nlp-45* in either the RMDD/V or RIA neurons in L1 stage animals (where normally only RIA expresses *nlp-45*) further reduced exploratory behavior below that of L1 controls **(Fig.5A)**. Altogether, these data suggest that *nlp-45* functions as an anti-exploratory neuropeptide.

**Figure 5:**
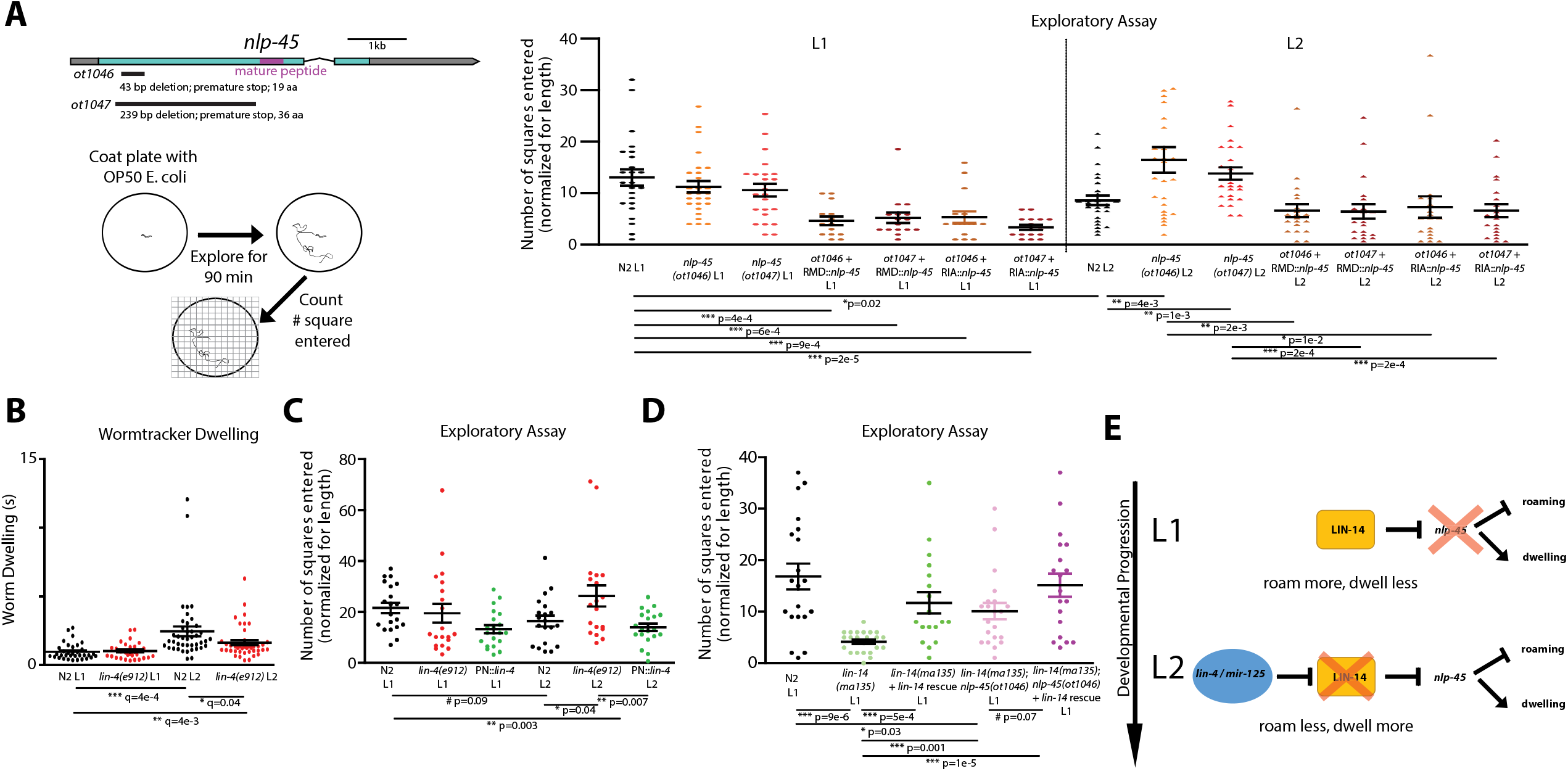
neuropeptide *nlp-45*, as controlled by *lin-4/lin-14*, regulates exploratory behavior across early larval transition. **(A)** *nlp-45* exerts temporally specific effects on anti-exploratory behavior. Schematic of *nlp-45* deletion mutants, generated by CRISPR-Cas9, is shown on the upper left panel: both *ot1046* and *ot1047* alleles results in frameshift mutations/premature stops and no production of the mature *nlp-45* peptide. On the bottom left is the schematic of the exploratory assay, as adapted and modified from Flavell *et al*., 2013. On the right is the quantification of the exploratory behavior, as measured by the number of squares entered normalized by the length of the animal. The mean +/− SEM are shown for each condition, and each point of the scatter dot plot represents a single animal. post hoc t-test *p<0.05; **p<0.01, ***p<0.001. **(B)** Increased dwelling during L1->L2 transition is partially juvenized by *lin-4(e912)* null mutation, as measured by automated worm tracking. The mean +/− SEM are shown for each condition, and each point of the scatter dot plot represents a single animal. Wilcoxon rank-sum tests and false-discovery rate q values for each comparison are shown below: *q<0.05; **q<0.01, ***q<0.001, **(C)** Neuronal *lin-4* regulates the “maturation” of exploratory behavior during L1->L2 transition. Exploratory behavior is measured as in (A). The quantification of the exploration behavior, as measured by the number of squares entered normalized by the length of the animal, is shown. PN::*lin-*4 represents transgenic strain *otTi71(Si[UPN::lin-4::unc54 3’utr])* with single copy insertion of panneuronal *lin-4* expression. The mean +/− SEM are shown for each condition, and each point of the scatter dot plot represents a single animal. post hoc t-test #0.05<p<0.1, *p<0.05; **p<0.01 **(D)** *lin-14(ma135)* null mutants exhibit “precocious” exploratory behavior through regulation of *nlp-45*. Exploratory behavior is measured as in (A). The mean +/− SEM are shown for each condition, and each point of the scatter dot plot represents a single animal. post hoc t-test #0.05<p>0.1, *p<0.05; ***p<0.001. **(E)** Schematic showing *lin-4*/*lin-14* regulation of *nlp-45* to alter exploratory behavior during the L1>L2 transition.

Next we tested whether the heterochronic regulators, *lin-4/mir-125* and its downstream target transcription factor *lin-14* are required for temporal transitions in exploratory behavior and, if so, whether they act through *nlp-45* to exert such effects. In two independent behavioral assays, we found that the L1->L2 behavioral transition in exploratory activity was partially juvenized (decreased dwelling/increase exploration) in *lin-4*/*mir-125* mutant (**Fig.5B, C**), which matched the juvenization of *nlp-45* expression pattern in *lin-4*/*mir-125* mutant animals. Conversely, forced neuronal expression of *lin-4*/*mir-125* at the L1 stage caused a precocious decrease in exploratory behavior (**Fig.5C**). Consistent with precocious (increased) *nlp-45* expression at the L1 stage in *lin-14* null animals (**Fig.4C**), these animals significantly reduced their exploratory behavior, and this effect was rescued by re-supplying *lin-14* (**Fig.5D**). Importantly, the decreased exploratory behavior in *lin-14* null mutant was partially suppressed in a *nlp-45* mutant background (**Fig.5D**). Together, these findings demonstrate that the *lin-4*/*mir-125* and *lin-14* regulation of *nlp-45* expression during early larval transition (L1 -> L2 onwards) contributes to the difference in exploratory behavior observed during this transition (**Fig.5E**).

#### *nlp-45* mediates transition in exploratory behavior upon sexual maturation

Given the role of *nlp-45* as an anti-exploratory neuropeptide during the L1 to L2 developmental transition, we next examined two other notable transitions in *C. elegans* post-embryonic development with reported changes in exploratory behavior. The first example was the increased exploration drive of adult males but not adult hermaphrodites for mate searching over food, observed upon sexual maturation and mediated by the PDF-1 neuropeptide (Barrios et al., 2012; Lipton et al., 2004). When we examined the *nlp-45* reporter allele at the onset of adulthood, expression emerged in specific neuron types in a sexually dimorphic manner. Specifically, in adult males, *nlp-45* was activated in 5 classes of neurons in the head (SAAD/V, AVA, RIV, AIB, RIM), most RVG and VA/VB/DA/DB/AS motor neurons in the VNC, as well as a number of tail neurons (**Fig.6A**). Also, expression of the reporter allele in all other neuron classes seemed to be generally elevated in adult males. Given the upregulation of *nlp-45* expression upon sexual maturation in adult males in comparison to adult hermaphrodites (**Fig.6A**), we investigated the role of *nlp-45* in the previously characterized food leaving behavior and found that *nlp-45* mutant adult males left food faster/earlier compared to control adult males (**Fig.6B**). *nlp-45* mutants did not significantly affect the food leaving behavior of juvenile males or adult hermaphrodites (**Fig.6B**). This suggested that the upregulation of *nlp-45* in adult males likely served as a counterbalancing anti-exploratory signal to the increased exploration drive as mediated by PDF-1.

**Figure 6:**
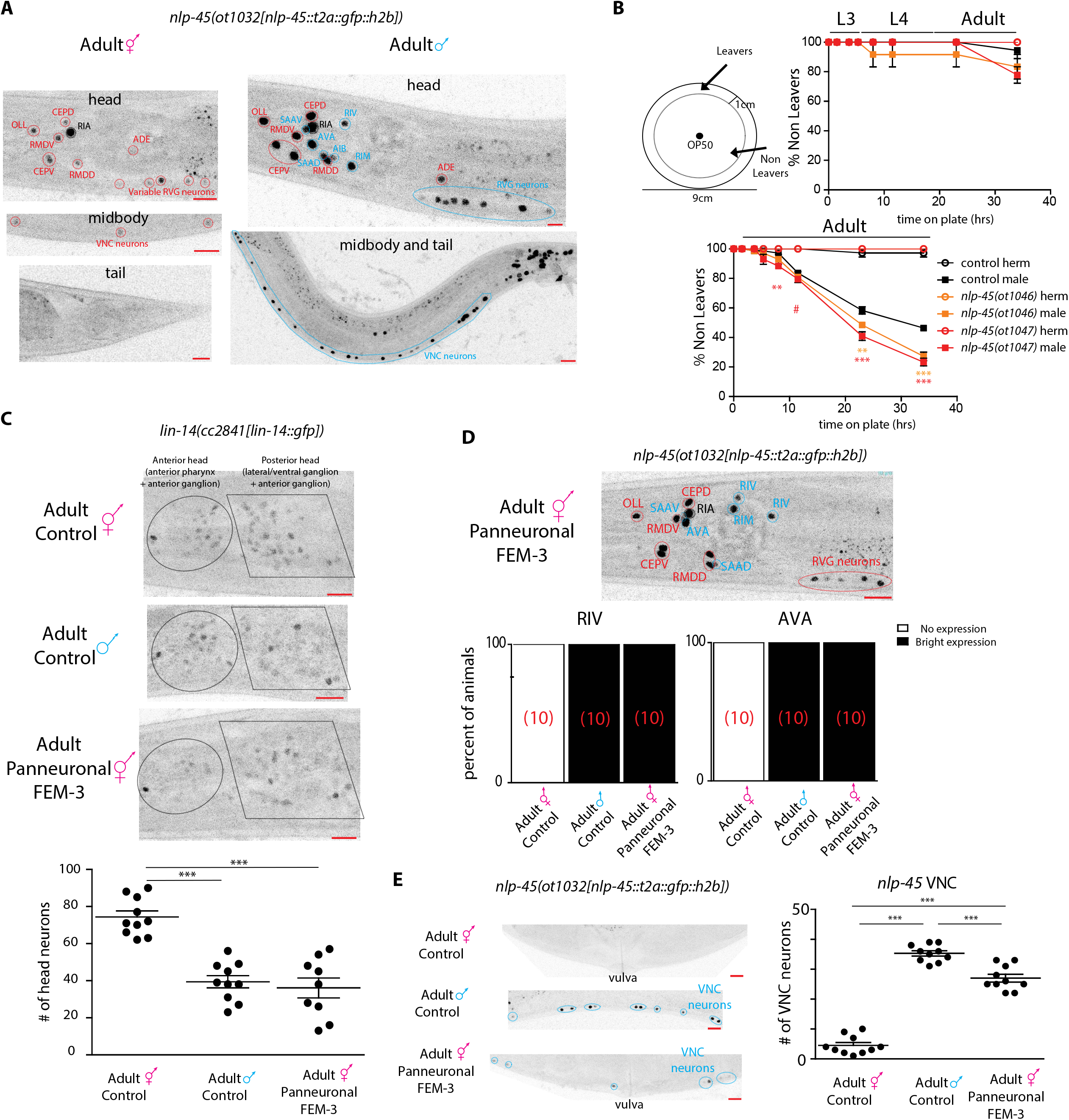
Sexually dimorphic regulation of *nlp-45*. **(A)** Expression of *nlp-45* in adult hermaphrodites and males. In addition to the neurons expressed in the later larval stages/adult hermaphrodite (labelled in red), those that are additionally turned on in the adult males are labelled in blue. Adult hermaphrodite head and midbody images are re-used from Fig.4A. **(B)** *nlp-45* promotes anti-exploratory behavior in adult males. Schematic of the leaving assay is shown on the upper left, as adapted from Lipton *et al*., 2004. On the upper right, *nlp-45* deletion mutants does not significantly affect the food leaving behaviors of juvenile males/hermaphrodites. On the bottom, *nlp-45* mutants specifically increase the food leaving behavior of adult males but not adult hermaphrodites. Values were plotted as mean +/− SEM of three independent experiments for juvenile animals and adult hermaphrodites and six independent experiments for adult males (n=12-16 animals per independent experiments). Statistical analysis is only shown for the comparison of *nlp-45* mutant adult males to control males (orange: *nlp-45(ot1046)* vs N2 adult males; red: *nlp-45(ot1047)* vs N2 adult males). post hoc t-test #0.05<p>0.1, *p<0.05; ***p<0.001 **(C)** Panneuronal depletion of sex determination master regulator TRA-1, through overexpression of FEM-3, decreases nervous system LIN-14 expression in adult hermaphrodites to mimic that of adult males. Representative microscope images, shown above, are overexposed in comparison to previous *lin-14* reporter images to better show the dim expression in adult males and FEM-3 overexpressed hermaphrodites. The quantifications of head neuron numbers across the three conditions are shown below. The mean +/− SEM are shown for each condition, and each point of the scatter dot plot represents a single animal. Post hoc t-test ***p<0.001. **(D)** Panneuronal depletion of sex determination master regulator TRA-1, through overexpression of FEM-3, masculinizes *nlp-45* expression in adult hermaphrodite head. Representative microscope image is shown above: additional *nlp-45* expression in SAAD/V, AVA, RIV, RIM neurons in FEM-3 overexpressed hermaphrodite, as compared to control hermaphrodite in panel A. Binary quantifications of *nlp-45* expression in the RIV and AVA neurons are shown below (number of animals for each condition is shown in red brackets). **(E)** Panneuronal depletion of TRA-1, through overexpression of FEM-3, masculinizes *nlp-45* expression in adult hermaphrodite VNC. Representative images are shown on the left. Quantifications of VNC neuron numbers are shown on the right. The mean +/− SEM are shown for each condition, and each point of the scatter dot plot represents a single animal. Post hoc t-test ***p<0.001. Red scale bars (10μm) are on the bottom right of all representative images.

We next examined the regulatory mechanism for *nlp-45*’s sexually dimorphic expression pattern. Intriguingly, *nlp-45* expression of the early larval *lin-14(0)* hermaphrodites largely mimicked that of the wild type adult males, with expression in neurons that was only observed in control adult males (e.g. SAAD/V, RIV, AIB, RIM) and also stronger expression in other neuron classes (e.g. OLL, RMDD/V, CEPD/V) (**Fig.4C, 6A**). This observation prompted us to ask whether the LIN-14 transcription factor is expressed in a sexually dimorphic manner. We indeed found that while LIN-14 expression was similarly downregulated in both sexes at early larval stages (L1-L3), its expression in the late larval (L4) and particularly in the adult stage was significantly more reduced in the male nervous system compared to that of the hermaphrodite (**Fig.6C, S3**). Altogether, this data indicated that the LIN-14 transcription factor maintained a juvenile *nlp-45* expression pattern by repressing it in specific neuron classes. The further downregulation of *lin-14* in the adult male nervous system allowed for the de-repression of *nlp-45* in additional neurons such as SAAD/V, RIV, AIB and RIM.

Since ChIP-seq analysis revealed binding of the hermaphrodite enriched master regulator of sexual identity, TRA-1, to *cis*-regulatory regions of *lin-14* (Berkseth et al., 2013), we examined the effect of TRA-1 on *lin-14* expression. To this end, we eliminated TRA-1 from the nervous system through panneuronal overexpression of FEM-3, a negative regulator of TRA-1 expression, frequently used to change the sexual identity of specific cell types (Lee and Portman, 2007; Oren-Suissa et al., 2016). We found that in these animals LIN-14 expression was significantly reduced in adult hermaphrodites and there was a consequent masculinization of *nlp-45* expression (e.g. expression in SAAD/V, AVA, RIV, RIM and VNC MNs) (**Fig.6C-E**). In summary, hermaphrodite enriched TRA-1 expression appears to maintain higher neuronal LIN-14 expression in adult hermaphrodites compared to adult males in order to prevent the onset of male-specific expression in specific neuron classes.

#### *nlp-45* mediates transition in exploratory behavior during the temporally and environmentally controlled dauer stage

The other notable example of transition in exploratory behavior was the previously observed increased locomotor quiescence upon entry into an alternative developmental stage, the diapause dauer stage (Bhattacharya et al., 2019; Cassada and Russell, 1975; Gaglia and Kenyon, 2009). The dauer stage is entered in response to harsh environmental conditions that are assessed at a precise developmental time point (Schaedel et al., 2012). In dauer animals, *nlp-45* gained expression in 6 classes (RMED/V, RMEL/R, IL1D/V, SAAV/D, RIV, RIM) of head neurons (**Fig.7A**). Investigating *nlp-45*’s role in the altered exploratory behavior in dauer animals, we found that *nlp-45* mutant dauer animals had reduced dwelling behavior compared to control dauer animals (**Fig.7B**). Hence, consistent with its anti-exploratory role in early larval stage transitions and upon sexual maturation in adult males, *nlp-45* upregulation in dauer animals contributes to the increased locomotor quiescence observed for this stage.

**Figure 7.**
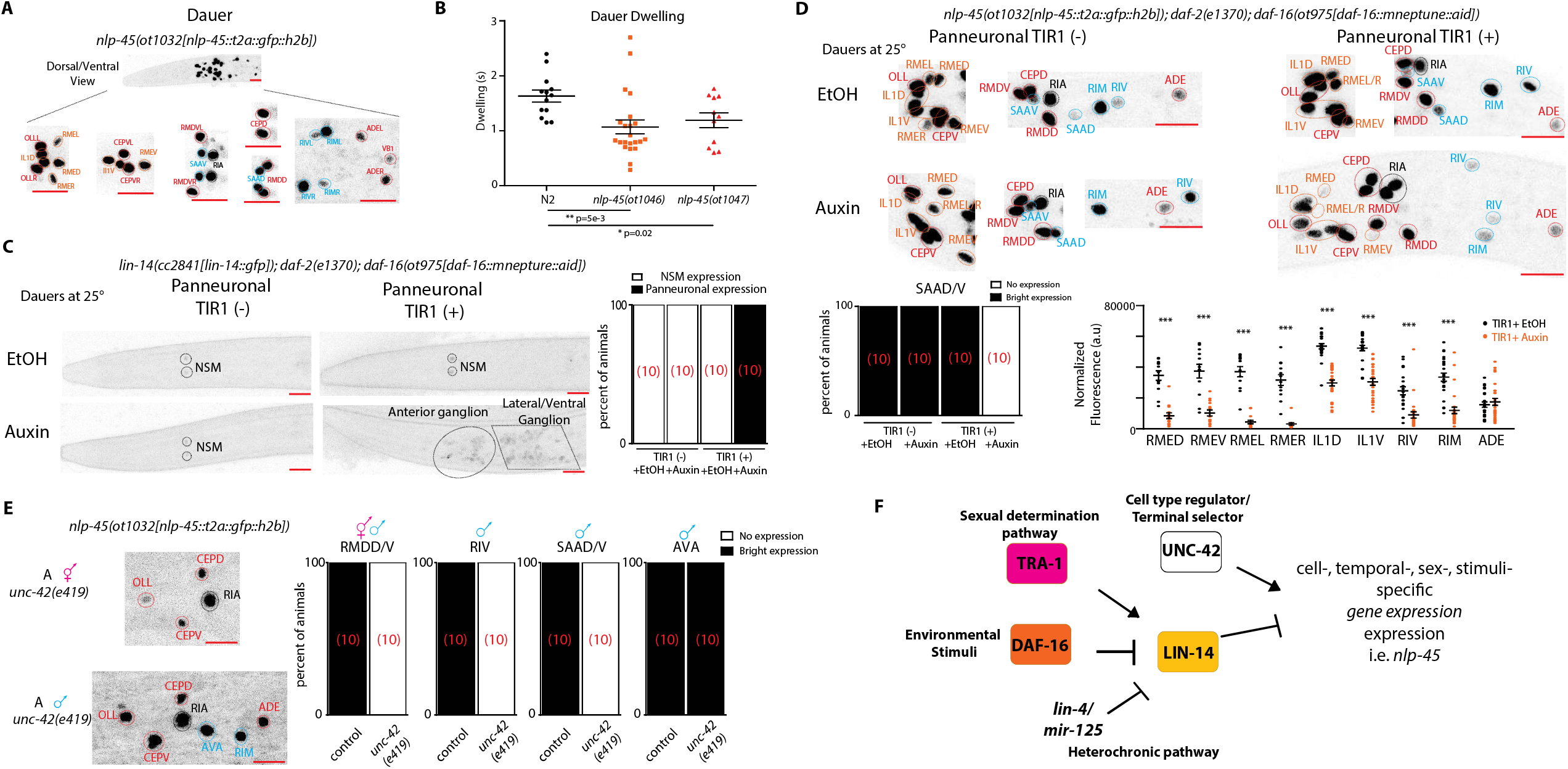
Intersectional control strategy dictates spatiotemporal expression pattern of *nlp-45*. **(A)** Expression of *nlp-45* in dauer animals. Upon entry into dauer, in addition to gaining expression in a number of neurons that are expressed in the adult male (labelled in blue), additional dauer specific expression is gained in a number of neurons (labelled in orange). For comparison to control mid-larval stages, see Fig. 4A. **(B)** *nlp-45* contributes to increased quiescence/dwelling behavior of dauer animals. *nlp-45* deletion mutant dauers exhibits reduced dwelling behavior compared to N2 dauers, as measured by automated worm tracker. The mean +/− SEM are shown for each condition, and each point of the scatter dot plot represents a single animal. post hoc t-test *p<0.05; **p<0.01 **(C)** Panneuronal degradation of DAF-16 in auxin-treated dauers leads to panneuronal de-repression of *lin-14* in dauer animals. Representative images are on the left while binary quantifications of panneuronal expression in are shown on the right (number of animals for each condition is shown in red brackets). **(D)** Panneuronal degradation of DAF-16 in auxin-treated dauers leads to a loss or reduced *nlp-45* expression in several neuronal classes. Representative images are on the top while quantifications are on the bottom. Binary quantifications are shown for the SAAD/V neurons (number of animals for each condition is shown in red brackets) while fluorescence quantifications are shown for the RMED/V, RMEL/R, IL1D/V, RIV, RIM, and ADE neurons. The mean +/− SEM are shown for each condition, and each point of the scatter dot plot represents a single animal. Post hoc t-test ***p<0.001. **(E)** Regulation of *nlp-45* by hardwired cell specific regulator, *unc-42*. Representative images of adult *unc-42(e419)* hermaphrodite and male animals are shown on the left while binary quantifications of *nlp-45* expression in the RMDD/V, RIV, SAAD/V and AVA neurons are shown on the right (number of animals for each condition is shown in red brackets). **(F)** Schematic showing temporal, sexual and environmental regulation of heterochronic regulator *lin-14*, in combination with cell specific terminal selector, *unc-42*, to dictate spatiotemporal gene batteries, e.g. *nlp-45*. Red scale bars (10μm) are on the bottom right of all representative images.

We next examined the regulatory mechanism for the temporally controlled transition of *nlp-45* expression in the dauer stage. Intriguingly, *nlp-45* expression in *lin-14(0)* hermaphrodites was observed in neurons that only showed *nlp-45* expression at the dauer stage (e.g. RMEs, IL1s) (**Fig.4C, 7A**). This suggested that, similarly to the sexually dimorphic regulation of *nlp-45*, environmental regulation of *nlp-45* expression may also converge on the regulation of transcription factor *lin-14*. Indeed, we found that this gain in *nlp-45* expression correlated with a global downregulation in LIN-14 in the dauer animals (**Fig.7C)**. Moreover, we found that this dauer-specific LIN-14 downregulation was cell-autonomously controlled by insulin-signaling in the nervous system (**Fig.7C**). Specifically, using an auxin-inducible degron system (Zhang et al., 2015), we observed that panneuronal depletion of a *daf-16/FoxO*, the key effector of insulin signaling, resulted in a specific de-repression of *lin-14* expression in the neurons of dauer stage animals (**Fig.7C**). This effect may be direct since ChIP-seq analysis revealed multiple DAF-16 *in vivo* binding sites in the *lin-14 cis*-regulatory region (Kumar et al., 2015). The de-repression of *lin-14* expression observed after neuronal *daf-16/FoxO* depletion led to the elimination/downregulation of *nlp-45* expression in the 6 classes of head neurons that gained expression upon entry into dauer (**Fig.7D**).

#### Intersection of plastic and hardwired control of *nlp-45* expression

To assess how the global temporal signal of LIN-14 activity results in highly neuron type-specific modulation of *nlp-45* expression, we turned to neuron type-specific transcription factors, terminal selectors, that specify and maintain neuron type-specific batteries of terminal identity genes (Hobert, 2016). We focused on the homeobox gene *unc-42/PROP1* which controls the differentiation of many *nlp-45* expressing neurons in both hermaphrodite and male animals (Baran et al., 1999; Pereira et al., 2015)(Berghoff et al., in preparation). We found that in *unc-42* mutant animals, *nlp-45* expression was affected in all neurons where *unc-42* and *nlp-45* expression normally overlap (e.g. RMDD/V, RIV, SAAD/V) with the exception of AVA, where *unc-42* only regulates a subset of its identity features (Baran et al., 1999; Pereira et al., 2015) (**Fig.7E**). We concluded that *unc-42* (and other terminal selector in other neurons) acts permissively to promote *nlp-45* expression, but that temporal, sexual and dauer signals mediated by *lin-14* antagonize the ability of *unc-42* to promote *nlp-45 e*xpression (**Fig. 7F**).

## DISCUSSION

Previous studies in both invertebrates and vertebrates have characterized many anatomical and electrophysiological maturation events in the postembryonic nervous system (Faumont et al., 2006; Gogtay et al., 2004; Okaty et al., 2009; White et al., 1978; Witvliet et al., 2020). Molecular changes that underlie these maturation events, as well as their genetic specification programs, have been less well analyzed. We presented here a comprehensive, nervous system-wide map of molecular changes that accompany the many behavioral transitions associated with post-embryonic nervous system maturation from juvenile to adult stages. Temporal transitions in gene expression were prominently observed for molecules with presumptive functions in neuronal communication. These included neurotransmitter receptors and gap junction molecules, consistent with vertebrate reports of altered chemical and electrical neurotransmission during post-natal development (Laurie et al., 1992; Nadarajah et al., 1997). Perhaps the most striking changes were observed in the neuropeptidergic system, mostly on the level of neuropeptide-encoding genes (**Fig.2, S1**). The pervasive expression change in the neuropeptide family across post-embryonic development, combined with anectodal evidence of dynamic neuropeptide expression controlling juvenile to adult transitions across many organisms (Conzelmann et al., 2013; Ellison et al., 2012; Lee et al., 2008; Schoofs and Beets, 2013; Wu et al., 2003), suggest that altering the repertoire of neuromodulatory peptides could be a conserved maturation mechanism in the animal kingdom that define behavioral state transitions across development. Specifically, we demonstrated that the spatiotemporal regulation of a novel anti-exploratory *nlp-45* across development resulted in consequential change in exploratory behavior across three separate temporal transitions.

Moving beyond the description of molecular changes, we characterized a genetic program that controls the temporal transitions in neuronal gene expression profiles across post-embryonic development. While studies in vertebrates have mostly focused on the role of environmental stimuli (e.g. neuronal activity) on the maturation of post-mitotic neuronal features, first exemplified by the studies of Hubel and Wiesel (Ben-Ari, 2002; Spitzer, 2006; Stroud et al., 2020; Wiesel and Hubel, 1963), much less attention had been focused on identifying the genetic programs that regulated the temporal identity of post-mitotic neurons, like those that have been characterized for the temporal identities of dividing neuroblasts (Cepko, 1999; Holguera and Desplan, 2018; Miyares and Lee, 2019; Pearson and Doe, 2004). Here, we identified heterochronic regulators, namely the microRNA *lin-4*, orthologous to *mir-125* in vertebrates, and its downstream target, the transcription factor *lin-14*, as a regulatory cassette that controls temporal transition in gene expression across post-embryonic development. *lin-4* and *lin-14* were first discovered and characterized for their role in controlling the temporal progression of mitotic cell types such as epithelial or reproductive cells in *C. elegans* (Rougvie and Moss, 2013). In the post-mitotic nervous system of *C. elegans*, some case studies in a few neuron types have characterized the role of *lin-4/lin-14* in controlling the temporal transitions in key neurodevelopmental events such as synaptic rewiring, axonal extension/branching, synaptogenesis and axonal degeneration (Hallam and Jin, 1998; Howell et al., 2015; Olsson-Carter and Slack, 2010; Ritchie et al., 2017; Xu and Quinn, 2016; Zou et al., 2012). Here, by profiling the adult neuronal transcriptome of *lin-4* and *lin-14* mutants, we demonstrated that they broadly regulated the temporal transitions in the expression of many but not all developmentally-regulated genes in the nervous system. We showed that the manipulation of *lin-4* and *lin-14* can juvenize behavioral patterns, and that the juvenized exploratory behavioral patterns are mediated by a *lin-4/lin14* target, the neuropeptide *nlp-45*. The role of *lin-4* in controlling neuronal maturation is conserved across species, as *mir-125*, the *lin-4* ortholog, has been shown to control several aspects of neuronal maturation in *Drosophila* and mice (Akerblom et al., 2014; Caygill and Johnston, 2008; Wu et al., 2012). In humans, duplication or deletion of chromosome 21 regions that include *mir-125* could result in intellectual and cognitive disabilities, a phenotype that could be considered a maturation defect (Elton et al., 2010; Errichiello et al., 2016). Another microRNA, *mir-101* has previously been shown to regulate circuit activity maturation (Lippi et al., 2016), indicating that microRNAs may be broadly employed in the control neuronal maturation. Moreover, the regulation of neuronal activity maturation by *mir-101* suggests a potential interplay between intrinsic genetic programs and neuronal activity in the regulation of neuronal maturation.

The observation that the sex determination pathway (through the global sex regulator TRA-1) and insulin responsive pathway (through DAF-16/FoxO) regulate LIN-14 expression suggest a complex, layered model of spatiotemporal gene regulation throughout post-embryonic nervous system development. Direct *in vivo* binding of TRA-1 and DAF-16 on *lin-14* were shown through ChIP analysis (Berkseth et al., 2013; Kumar et al., 2015); here we demonstrated that TRA-1 and DAF-16 are indeed required for the dynamic control of *lin-14* expression. The modulation of the heterochronic regulators by these other pathways suggests that a significant subset of temporally regulated *lin-14* targets are likely to be sexually dimorphic and plastic in response to environmental stress, as observed for *nlp-45*. The observation that these temporally regulated genes are under the control of terminal selector regulation suggests a perhaps generalizable model of spatiotemporal gene expression regulation throughout post-embryonic development: Cell-specific terminal selectors specify the spatial identity of each neuron’s gene battery consisting of a subset of static genes, that maintain the same expression throughout the lifetime of the organism, and a subset of temporally regulated genes. Temporal regulation may be imposed on terminal selector-controlled genes by heterochronic regulators either promoting or antagonizing the ability of terminal selectors to activate their target genes.

## ACKNOWLEDGMENTS

We thank Qi Chen for generating transgenic lines, Dylan Rahe for help with INTACT optimization, Alex Romero and Eviatar Yemini for help with worm tracking, Abhishek Bhattacharya for integration of the *inx-19* fosmid reporter, Berta Vidal Iglesias for integration of the *nlp-13* promoter fusion reporter, Luisa Cochella, Michael P Hart, and Isabel Beets for comments on the manuscript, members of the Hobert lab for resources and comments on the manuscript. This work was funded by the NIH K99 HD098371, National Research Council of Canada (Holmes Award), and by the Howard Hughes Medical Institute. Some strains were provided by the CGC, which is funded by NIH Office of Research Infrastructure Program (P40 OLD010440).

## AUTHOR CONTRIBUTIONS

Conceptualization, H.S. and O.H.; Methodology, H.S.; Validation, H.S.; Formal Analysis, H.S.; Investigation, H.S.; Resources, H.S.; Writing – Original Draft, H.S.; Writing – Review & Editing, H.S. and O.H.; Visualization, H.S.; Supervision, H.S. and O.H; Funding Acquisition, H.S. and O.H.

---

**Figure S1:**
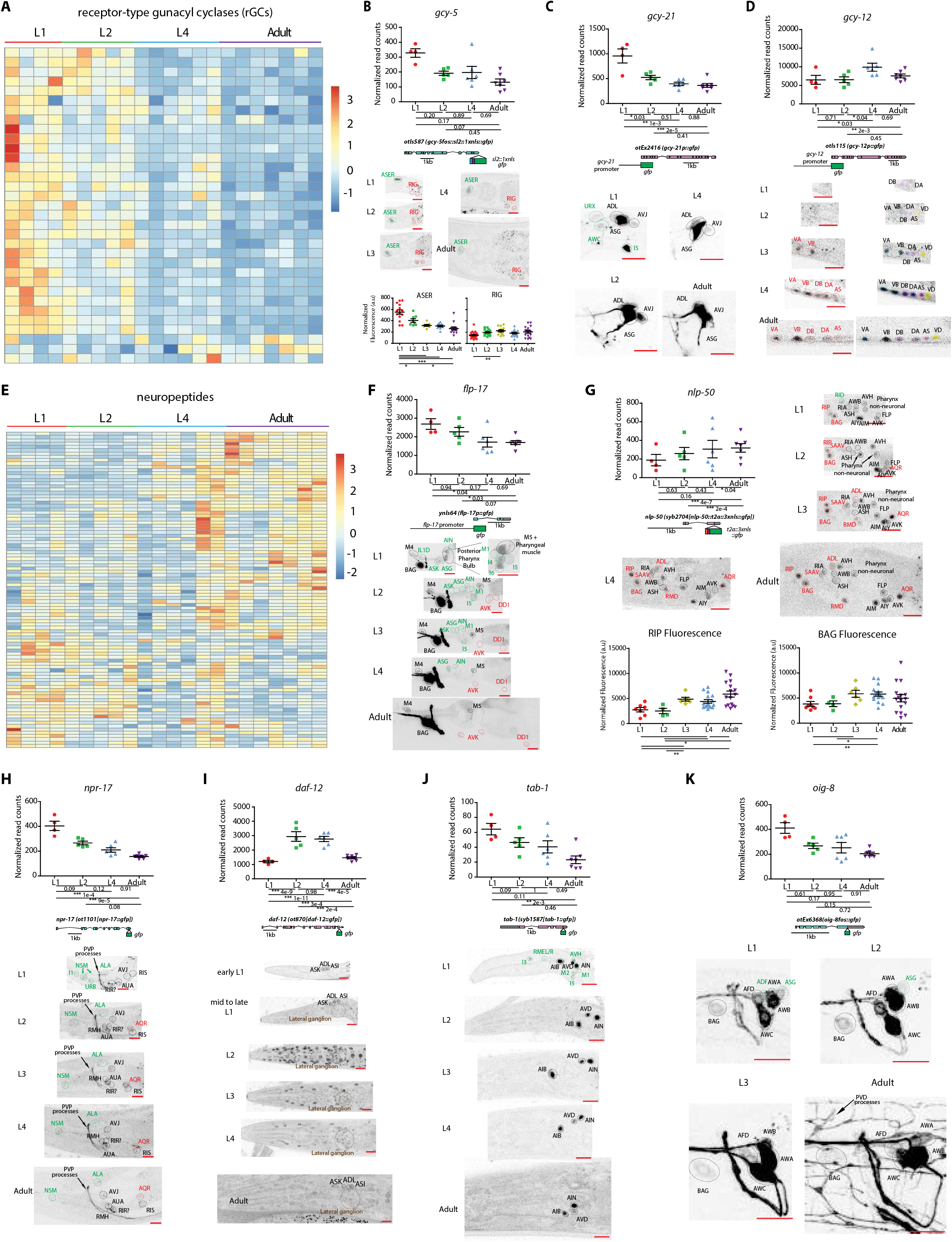
Additional validation of nervous system gene expression changes across *C. elegans* post-embryonic development. **(A)/(E)** Heatmap of all neuronally enriched receptor-type gunacyl cyclases (rGCs) (A) and neuropeptides (E) across post-embryonic development. Values were z-score normalized and plotted using pheatmap in R studio. Each row represents a single gene, and each column represents a single RNA-seq replicate. For (B)-(D), (F)-(K), validations of developmentally regulated genes with expression reporters are shown. On top are the scattered dot plots (each point represents a single replicate) of the normalized read counts across all developmental stages from the neuronal INTACT/RNA-seq profiling. Mean +/− SEM are shown for each stage. Adjusted p values (p_adj_) for each developmental comparison are below. For p_adj_, *<0.05, **<0.01, ***<0.001. Below the RNA-seq read count plots are the schematics and allele names of the expression reporters. Below that are representative confocal microscopy images of the expression reporters across development. Specific regions/neurons labeled with dotted lines: those labelled with black dotted lines/names are not altered developmentally while those labelled with green and red lines/names demonstrate, respectively, decreases and increases in expression across development. Those labelled with brown lines/names demonstrate both increases and decreases in expression in the same neurons across development. For (B) and (G), additional quantifications of fluorescence intensity are also shown. Red scale bars (10μm) are on the bottom right of all representative images. For all panels, L1 through L4 represent the first through the fourth larval stage animals. **(B)** rGC *gcy-5*, as validated with a transcriptional fosmid reporter (*sl2::1xnls::gfp*), shows decreased expression in ASER and increased expression in RIG, as measured with fluorescence intensity, across development. **(C)** rGC *gcy-21*, as validated with a promoter fusion reporter (*gfp*), loses expression in three neuronal classes across the L1->L2 transition. **(D)** rGC *gcy-12*, as validated with a promoter fusion reporter (*gfp*), gains expression in A and B type motorneurons across mid/late larval development. **(F)** Neuropeptide-encoding gene *flp-17*, as validated with a promoter fusion reporter (*gfp*), loses and gains expression in nine and two classes of neurons, respectively. **(G)** Neuropeptide-encoding gene *nlp-50*, as validated with an endogenous reporter (*t2a::3xnls::gfp*) engineered with CRISPR-Cas9, gains expression in in a number of neurons across development. It also loses expression in the RID neuron during early larval development. **(H)** Neuropeptide receptor gene *npr-17*, as validated with an endogenous translational reporter (*gfp*) engineered with CRISPR-Cas9, shows decreased and increased expression in four and one classes of neurons, respectively. **(I)** Nuclear hormone receptor *daf-12*, as validated with an endogenous translational reporter (*gfp*) engineered with CRISPR-Cas9, shows increased expression broadly across the nervous system during early/mid-larval stage and then decreased expression upon transition into late larval/adult stage. **(J)** Homeodoman transcription factor *tab-1*, as validated with an endogenous translational reporter (*gfp*) engineered with CRISPR-Cas9, loses expression in five classes of neuron during early larval development. **(K)** Immunoglobulin-like domain molecule *oig-8*, as validated with a translational fosmid reporter (*gfp*), loses expression in two classes of neuron during early larval development.

**Figure S2:**
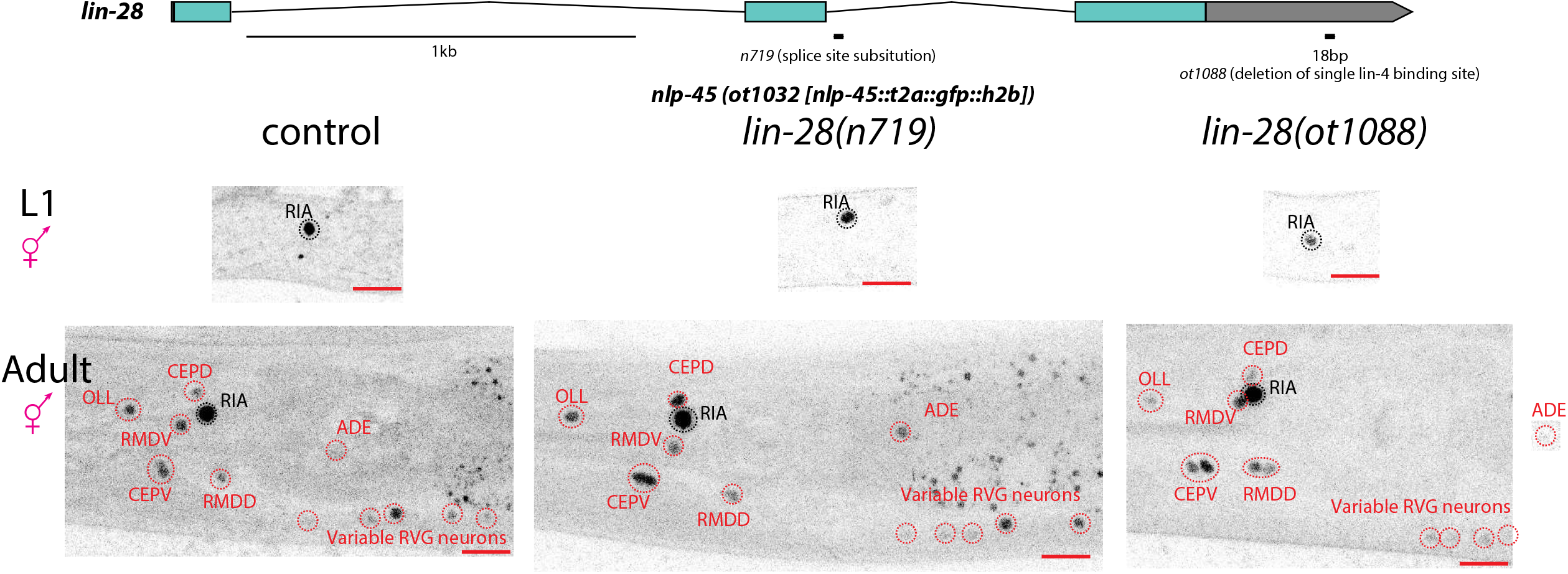
*lin-28* does not regulate temporal expression pattern of *nlp-45*. Schematic of *lin-28(n719)* loss of function and *lin-28(ot1088)* gain of function alleles is shown above. Shown below are representative images of the *nlp-45* expression reporter in control, *lin-28(n719)* and *lin-28(ot1088)* animals. Neurons that are labelled in black are not developmentally regulated while those that are labelled in red are developmentally upregulated. Red scale bars (10μm) are on the bottom right of all representative images.

**Figure S3:**
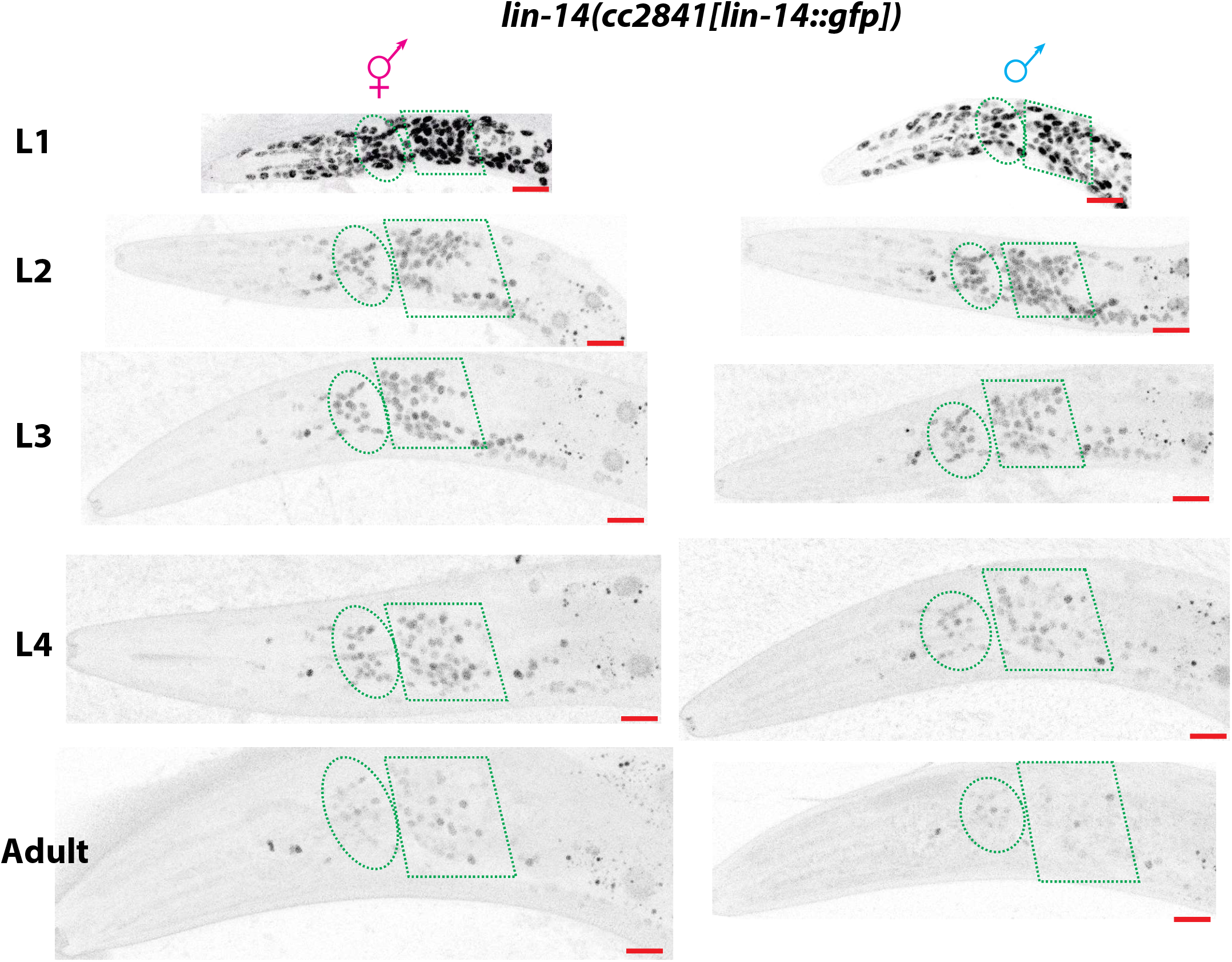
Developmental expression of *lin-14* in hermaphrodite and male animals. Representative images for the *lin-14(cc2841[lin-14::gfp])* reporter, as engineered by CRISPR-Cas9, is shown across all developmental stage for hermaphrodite and male animals. Ellipse and polygon outlines the anterior and lateral/ventral neuronal ganglions respectively. Representative images for hermaphrodite are re-used here from Fig. 3C for direct side by side comparison with male animals across development. Red scale bars (10μm) are on the bottom right of all representative images.

## METHODS

### *C. elegans* strains and handling

Worms were grown at 20°C on nematode growth media (NGM) plates seeded with E. coli (OP50) bacteria as a food source unless otherwise mentioned. Worms were maintained according to standard protocol. Wild-type strain used is Bristol variety, strain N2. A complete list of strains and transgenes used in this study is listed in the Table S6. Whenever synchronization of developmental stage was necessary, animals were egg prepped according to standard protocol and synchronized at the L1 stage. They were then plated on food and collected after 8 +/−1 hrs, 21 +/−1 hrs, 30 +/− 2hr, 40 +/− 2hrs, and 53 +/− 2hrs for L1, L2, L3, L4, and adult stages, respectively for either molecular or behavioral analysis. These time points were chosen such that the animals were in the middle of each larval stage or relatively early in adulthood for the analysis. Dauer animals were obtained using standard crowding, starvation and high temperature conditions.

### Constructs cloning and stain generation

#### UPN INTACT

To generate the UPN::INTACT tag (*npp-9::mcherry::3xflag*), a concatenated panneuronal promoter (Yemini et al., 2019), containing promoter fragments from *unc-11, rgef-1*, *ehs-1*, and *ric-19*, and the INTACT tag (Steiner et al., 2012) were cloned together using Gibson assembly. The constructs was injected (5ng/μl with 100ng/ μl digested OP50 DNA) and the resulting extrachromosomal array strain was integrated into the genome using standard UV irradiation methods. This was followed by 6 rounds of backcrossing to N2 to generate *otIs790*.

#### Fosmid recombineering

To generate the *lin-4* fosmid reporter, standard fosmid recombineering protocol was used as described previously (Tursun et al., 2009). Briefly, a 90bp *lin-4* primary miRNA was replaced with *nls::yfp::h2b* in fosmid WRM0613aD08. The recombineered fosmid was injected (15ng/μl with 100ng/μl digested OP50 DNA), and the resulting extrachromosomal array strain was integrated into the genome using standard UV irradiation methods. This was followed by 2 rounds of backcrossing to N2 to generate *otIs763*.

#### Genome engineering (CRISPR-Cas9)

*lin-14(ot1087), lin-28(ot1088), nlp-45(ot1032[nlp-45::t2a::gfp::h2b), nlp-45(ot1046), nlp-45(ot1047)* were generated using Cas9 protein, tracrRNA, and crRNAs from IDT, as previously described (Dokshin et al., 2018). For *lin-14(ot1087)*, two crRNAs (gttcctgagagcaatttttg and caaaactcacaaccaactca) and a single strand oligodeoxynucleotide (ssODN) donor (ttgctttttcctgcactcactttacctttgtctcactttttcttacttctgtatcacaaaaatgattata) was used to ensure a precise 466bp deletion in the *lin-14* 3’UTR to remove all seven *lin-4* binding site. For *lin-28(ot1088)*, one crRNA (cctgagagtgcaatttgagg) and a ssODN donor (cccctctaaaccatactaccacctacctcctcaaacttttttttttcaaatagaactgattgcacctgtt) were used to ensure a precise 18bp deletion in the *lin-28* 3’UTR to remove the single *lin-4* binding site. For, *nlp-45(ot1032[nlp-45::t2a::gfp::h2b)*, two crRNAs (aagcatctggactgccgatg and tgacttgaacaggaagcatc) and an asymmetric double stranded *t2a::gfp::h2b*, PCRed from pBALU43 (Tursun et al.), were used to insert the fluorescent tag at the C terminal. For *nlp-45(ot1046)* and *nlp-45 (ot1047*), two crRNAs (acttgcgttaaccacaatga and tgacttgaacaggaagcatc) were used and random deletions were screened to obtain *nlp-45(ot1046)* and *nlp-45 (ot1047)*. These deletions were 43 and 239 bps within the first exon, respectively, and both resulted frameshift mutations and premature stops. Neither mutation resulted in the production of the predicted mature peptide (Fig. 6C). *npr-17(ot1101)* was generated using standard method as previously described to insert C-terminal fused *gfp* (Dickinson et al., 2015). *ins-6(syb2685)*, *ins-9 (syb2616)*, *nlp-50(syb2704)* were generated by SUNY Biotech. To facilitated the neuronal ID of the secreted neuropeptide expression reporters, a nuclear localized GFP was inserted behind the neuropeptide coding sequences, separated by a T2A sequence that splits the two proteins (Ahier and Jarriault, 2014).

#### Single copy insertion by MiniMos

The concatenated panneuronal promoter (UPN) and a 338bp fragment containing the *lin-4* miRNA were fused together and cloned into pCFJ910 using Gibson Assembly. The plasmid was injected to obtain single copy insertion of UPN::*lin-4* as previously described (Frokjaer-Jensen et al., 2014).

#### Cell specific nlp-45 overexpression

*mgl-1* promoter (Zhang et al., 2014) and *glr-3* promoter (Brockie et al., 2001) were PCRed from genomic DNA for neuron specific expression in RMDD/V and RIA neurons respectively. *nlp-45* cDNA was obtained from Dharmacon. The promoter fragments and the *nlp-45* cDNA were fused together with *sl2::2xnls::tagrfp::p10 3’utr* by Gibson assembly. The constructs were injected at 50ng/μl and extrachromosomal array lines were selected according to standard protocol.

#### All other strains

The *inx-19* fosmid (*otIs773*) was obtained throughout integration of a previously published extrachromosomal array strain (Bhattacharya et al., 2019). The *nlp-13* promoter fusion GFP reporter (*otIs742*) was obtained through integration of an existing extrachromosomal array strain, HA329, and backcrossed 5x. The *gcy-12* promoter fusion GFP reporter was obtained through integration of an existing extrachromosomal array strain, DA1266. All other strains were previously published, and/or obtained from CGC and/or crosses with these strains as detailed in the Key Resources Table.

### Neuron identification

For neuronal cell identification, colocalization with the NeuroPAL landmark strain (*otIs669* or *otIs696*) was used to determine the identity of all neuronal expression as previously described (Yemini et al., 2019).

### Behavioral analysis

#### Automated Worm tracking

Automated single worm tracking was performed using the Wormtracker 2.0 system at room temperature (Yemini et al., 2013). Animals at all stages were recorded for 5 mins each except for dauers, which were recorded for 10 mins to ensure adequate sampling of locomotor features due to increased quiescence. All animals were tracked on NGM plates uniformly covered with food (OP50), except for dauers, which were tracker on non-coated plates. Analysis of the tracking videos was performed as previously described (Yemini et al., 2019).

#### Exploratory Assay

To measure exploration behavior, an adapted exploratory assay from a previous study (Flavell et al., 2013) was used to increase sensitivity for younger/smaller animals. Individual animals at the respective developmental stages and genotypes were picked to a 5 cm agar plate uniformly seeded with *E. coli* strain OP50. After 90 min, plates were superimposed on a grid containing 1 mm squares, and the number of squares entered by the worm tracks were manually counted. The number of squares explored is adjusted for the length of the animal, to compensate for the different size of animals at different developmental stages/genotypes. Transgenic and mutant strains were always compared to control animals assayed in parallel. All plates were scored by an investigator blind to the genotype of the animals.

#### Food leaving Assay

The food leaving (also known as mate-searching) assay was performed as previously described (Lipton et al., 2004). A single drop (18μl) of OP50 was seeded the day before and allowed to grow. The following day, a single animal of the respective developmental stage and sex was placed in the center of a 9 cm agar plate, and each animal that had left the food was scored blindly at 8 time points for a period of 34 hrs. A worm was considered a leaver if it was 1 cm from the edge of the plate.

### Microscopy

Worms were anesthetized using 100mM of sodium azide and mounted on 5% agarose on glass slides. All images were acquired using a Zeiss confocal microscope (LSM880). Image reconstructions was perform using Zen software tools. Maximum intensity projections of representative images were shown. Fluorescence intensity was quantified using the Zen software. Figures were prepared using Adobe Photoshop and Illustrator.

### INTACT for purification of affinity-tagged neuronal nuclei

UPN::INTACT control worms (*otIs790*) as well as mutants were grown on large plates (150mm) with enriched peptone media coated with NA22 bacteria to allow for the growth of large quantities of worms: 100,000 worms can grow from synchronized L1 stage to gravid adults on a single plate. Animals were collected at the respective stage as described above. *lin-4(e912)* and *lin-14(ot1087)* animals were slower in their developmental progression compared to controls (Chalfie et al., 1981), and adult *lin-4(e912)* and *lin-14(ot1087)* animals were collected ~ 57+/− 2 hrs (4 hrs after the control and *lin-28(ot1088)* adult animals were collected). ~600,000 animals were collected for each L1/L2 replicates, while ~200,000 animals were collected for each L4/adult replicates. At the time of collection, animals were washed off the plate with M9, washed 3x with M9, lightly fixed with cold RNAse-free DMF for 2 minutes before washing with 1xPBS 3x.

Modification were made from the previous INTACT protocol (Steiner et al., 2012) to optimize pulldown of neuronal nuclei. All steps following were done in cold rooms (4°C) to minimize RNA and protein tag degradation. The animals were homogenized mechanically using disposable tissue grinders (Fisher) in 1x hypotonic buffer (1x HB: 10mM Tris pH 7.5, 10mM NaCl, 10mM KCl, 2mM EDTA, 0.5mM EGTA, 0.5mM Spermidine, 0.2mM Spermine, 0.2mM DTT, 0.1% Triton X-100, 1x protease inhibitor). After each round of mechanical grinding (60 turns of the grinder), the grinder was washed with 1mL 1x HB and the entire homogenate was centrifuged at 100xg for 3 min. The supernatant was collected for later nuclei extraction and the pellet was put under mechanical grinding and centrifugation for 4 additional rounds. The supernatant collected from each round were pooled, dounced in a glass douncer, and gently passed through an 18-gauge needle 20x to further break down small clumps of cells. The supernatant was then centrifuged at 100xg for 10 min to further remove debris and large clumps of cells. Nuclei was isolated from the supernatant using Optiprep (Sigma): supernatant after centrifugation was collected in a 50mL tube, added with nuclei purification buffer (1x NPB: 10mM Tris pH 7.5, 40mM NaCl, 90mM KCl, 2mM EDTA, 0.5mM EGTA, 0.5mM Spermidine, 0.2mM Spermine, 0.2mM DTT, 0.1% Triton X-100, 1x protease inhibitor) to 20mL, and layered on top of 5mL of 100% Optiprep and 10mL of 40% Optiprep. The layered solution was centrifuged at 5000xg for 10 min in a swinging bucket centrifuge at 4°C. The nuclei fraction was collected at the 40/100% Optiprep interface. After removal of the top and bottom layers, leaving a small volume containing the nuclei, the process was repeated 2 additional times. After final collection of the crude nuclei fraction, the volume was added to 4mL with 1xNPB and precleared with 10μl of Protein-G Dynabeads and 10 μl of M270 Carboxylated beads for 30min to 1hr (Invitrogen). The precleared nuclei extract was then removed, and 50 μl was taken out as input samples (total nuclei). The rest was incubated with 30μl of Protein G Dynabeads and 3 μl of anti-FLAG M2 antibody (Sigma) overnight to immunoprecipitate (IP) the neuronal nuclei. The following day, the IPed neuronal nuclei/beads was washed 6-8 times with 1xNPB for 10-15 min each time. The resulting IPed neuronal nuclei/beads were resuspended in 50 ul 1xNPB and a small aliquot was used to check with DAPI staining to quality-check the procedure for the following: 1) sufficient quantities of nuclei was immunoprecipitated; 2) nuclei are intact and not broken; 3) the majority of bound nuclei are single, mCherry-labelled neuronal nuclei and minimal nuclei clumps and large tissue chunks were immunoprecipitated. Anything not satisfying these quality checks were not used for downstream processing. The resulting input and neuronal IP samples were used for isolation of total RNA using Nucleospin RNA XS kit according to manufacturer’s protocol (Takara).

### RNA-seq and data analysis

RNA-seq libraries were prepped using the Universal RNA-seq kit (Tecan) according to manufacter’s protocol. The libraries were sequenced on Illumina NextSeq 500 machines with 75bp single-end reads. After initial quality check, the reads were mapped to WS220 using the Subread package (Liao et al., 2019), and assigned to genes using featurecounts. Neuronal enrichment was conducted by comparing neuronal IP samples to their respective input samples using DESeq2, with batch effect taken into account for the analysis (Love et al., 2014). 7974 genes were found to be neuronally-enriched (**Table S1**). We took the read counts of these 7974 genes for all IP samples across development, normalized for library size, and conducted all developmental and mutant analysis using DESeq2. We found that this approach minimized contamination artifacts resulting from the protocol and led to the best biological validation.

### Auxin inducible degradation

The AID system was employed as previously described (Bhattacharya et al., 2019; Zhang et al., 2015). The conditional *daf-16* allele *daf-16(ot975[daf-16::mneptune2.5::3xflag::aid)* (Aghayeva et al., 2020) was crossed with *daf-2(e1370)*, panneuronal TIR1-expressing transgenic lines and *lin-14*/*nlp-45* reporters to generate the experimental strains. Animals were grown (from embryo onward) on NGM plates supplemented with OP50 and 4mM auxin in EtOH (indole-3 acetic acid, IAA, Alfa Aesar) at 25°C to degrade DAF-16 panneuronally and to induce dauer formation. As controls, plates were supplemented with the solvent EtOH instead of auxin. Additional control animals without panneuronal TIR-1 expression grown on EtOH and auxin were also included for comparison.

### QUANTIFICATION AND STATISTICAL ANALYSIS

Statistical analysis of the automated worm tracking videos was performed as previously described (Yemini et al., 2013). Briefly, statistical significance between each group were blindly calculated using Wilcoxon rank-sum test and correcting for false-discovery rate. Statistical analysis of RNA-seq comparison was performed using DESeq2 as previously described (Love et al., 2014). All microscopy fluorescence quantifications were done in the Zen software (Carl Zeiss). For all behavioral assay, randomization and blinding was done wherever possible. All statistical tests for fluorescence quantifications and behavior assays were conducted using Prism (Graphpad) as described in figure legends.

## SUPPLEMENTAL TABLE LEGENDS

**Table S1: Neuronally-enriched genes**

Comparison of neuronal-nuclei immunoprecipitated (IP) samples to input (total nuclei) samples is conducted using DESeq2 (Love et al., 2014) in R Studio to determine enrichment. 7974 genes have a log2FoldChange>0 (neuronally-enriched over input) and a p_adj_< 0.05. The genes are sorted by p_adj_ from smallest to largest. baseMean is the average normalized read counts of all IP and input samples. log2FoldChange is calculated using the formula log2 (average read counts of IP samples)/(average read counts of Input samples).

**Table S2: Normalized read counts of all 7974 neuronally-enriched genes across post-embryonic development**

Raw read count for the 7974 neuronally-enriched genes are extracted for the neuronal IP samples and adjusted for library size. These are then used in DESeq2 (Love et al., 2014) to conduct comparisons between developmental stages, as shown in Table S3-4. The final read counts displayed are as a result of normalization done in the DESeq2 program. There are 4, 5, 6, 7 replicates shown for L1, L2, L4, and adult stages respectively.

**Table S3: Comparison of all temporal transitions among the 7974 neuronally-enriched genes across post-embryonic development**.

Pairwise comparisons between all stages are conducted using DESeq2 (Love et al., 2014) as described in Table S2. The log2FoldChange and p_adj_ are shown for each comparison. All 7974 neuronally-enriched genes are shown regardless of p_adj_. The table is sorted alphabetically by gene name.

**Table S4: Normalized read counts of 2639 developmentally regulated genes**

Same as Table S2, but only with a subset of 2639 genes that have a p_adj_ <0.01 in any pairwise comparisons between developmental stages.

**Table S5: Genes that are juvenized in adult heterochronic mutants**

Pairwise comparisons between adult *lin-4(e912)* null, *lin-14(ot1087)* gain of function, and *lin-28(ot1088)* gain of function mutant neuronal IPs are conducted against adult control neuronal IP data, using only the read counts of the 7974 neuronally-enriched genes. The log2FoldChange and p_adj_ are shown for each comparison. All 7974 neuronally-enriched genes are shown regardless of p_adj_. The table is sorted alphabetically by gene name. log2FoldChange is calculated using the formula log2 (average read counts of mutant adult samples)/(average read counts of control adult samples).

**Table S6: Reagents and Strains Used in this manuscript**

